# Targeted Degradation of CDK9 Potently Disrupts the MYC Transcriptional Network

**DOI:** 10.1101/2024.05.14.593352

**Authors:** Mohammed A. Toure, Keisuke Motoyama, Yichen Xiang, Julie Urgiles, Florian Kabinger, Ann-Sophie Koglin, Ramya S. Iyer, Kaitlyn Gagnon, Amruth Kumar, Samuel Ojeda, Drew A. Harrison, Matthew G. Rees, Jennifer A. Roth, Christopher J. Ott, Richard Schiavoni, Charles A. Whittaker, Stuart S. Levine, Forest M. White, Eliezer Calo, Andre Richters, Angela N. Koehler

**Affiliations:** Koch Institute for Integrative Cancer Research, Massachusetts Institute of Technology, 500 Main Street, Cambridge, MA 02139, USA; Department of Biological Engineering, Massachusetts Institute of Technology, Cambridge, MA 02139, USA; MIT Center for Precision Cancer Medicine, Massachusetts Institute of Technology, Cambridge, MA 02139, USA; Broad Institute of MIT and Harvard, Cambridge, MA 04142, USA; Harvard-MIT Health Sciences and Technology, Boston, MA 02115, USA; Massachusetts General Hospital Cancer Center, Charlestown, MA 02129, USA; Department of Medicine, Harvard Medical School, Boston, MA 02115, USA; MIT BioMicro Center, Department of Biology, Massachusetts Institute of Technology Cambridge, MA, 02139 USA; Department of Biology, Massachusetts Institute of Technology, Cambridge, MA 02139, USA

## Abstract

Cyclin-dependent kinase 9 (CDK9) coordinates signaling events that regulate RNA polymerase II (Pol II) pause-release states. It is an important co-factor for transcription factors, such as MYC, that drive aberrant cell proliferation when their expression is deregulated. CDK9 modulation offers an approach for attenuating dysregulation in such transcriptional programs. As a result, numerous drug development campaigns to inhibit CDK9 kinase activity have been pursued. More recently, targeted degradation has emerged as an attractive approach. However, comprehensive evaluation of degradation versus inhibition is still critically needed to assess the biological contexts in which degradation might offer superior therapeutic benefits. We validated that CDK9 inhibition triggers a compensatory mechanism that dampens its effect on MYC expression and found that this feedback mechanism was absent when the kinase is degraded. Importantly, CDK9 degradation is more effective than its inhibition for disrupting MYC transcriptional regulatory circuitry likely through the abrogation of both enzymatic and scaffolding functions of CDK9.

**Highlights:**

– KI-CDK9d-32 is a highly potent and selective CDK9 degrader.
– KI-CDK9d-32 leads to rapid downregulation of MYC protein and mRNA transcripts levels.
– KI-CDK9d-32 represses canonical MYC pathways and leads to a destabilization of nucleolar homeostasis.
– Multidrug resistance ABCB1 gene emerged as the strongest resistance marker for the CDK9 PROTAC degrader.

## Introduction

*MYC* gene expression is an important hallmark of stimulated signaling pathways that promote cell proliferation^1,2^. Dysregulation of *MYC* expression resulting from genomic amplification or increased gene copy number of the gene, among a variety of other genomic alterations, is a key driver in cancer development and progression^3^. Thus, the suppression of MYC transcription and its downstream programs has been a long-standing goal in the development of cancer therapeutics.

CDK9 is the catalytic subunit of the positive transcription elongation factor b (P-TEFb), and is crucial for the upstream and downstream regulation of MYC. MYC gene expression depends highly on CDK9 which is tethered to the MYC promoter through the scaffolding activity of the ERK protein^1^. Moreover, CDK9’‘s influence on MYC dynamics extends to the protein level as well. CDK9 mediated phosphorylation of MYC at serine 62 protects the oncoprotein from degradation^4^. Consequently, MYC genomic amplification induces an increased dependence on CDK9 which in turn becomes critical for the maintenance of the MYC addicted tumor state^5^. More than twenty CDK9 inhibitors have been evaluated in clinical trials involving both hematologic and solid tumors. Unfortunately, a combination of off- and on-target toxicity and the lack of objective response has restricted progress to FDA approval for these agents^6^. Sustained inhibition of CDK9 can induce a compensatory increase in *MYC* expression^2^. This mechanism of resistance is driven by activation of inactive cellular CDK9 through the bromodomain protein BRD4 and its channeling of available CDK9 to the MYC promoter^2^.

The sustained interest in advancing molecules with desirable pharmacological attributes against MYC and CDK9 has been bolstered by advances in targeted protein degradation^7^. It is likely that an acute and potent degradation of CDK9 would circumvent the resistance mechanism described above and lead to a more robust attenuation of MYC activity. While the pharmacodynamics of inhibitors are driven by drug concentration, those of degraders appear to be driven by the target resynthesis rate^8^. Moreover, degraders have been shown to act sub-stoichiometrically with rapid kinetics^7^. These attributes may confer PROTACs and other degraders an advantage in the form of greater resilience to drug resistance induced by long-term exposure to high concentration of small molecule inhibitors^7,9^.

Moreover, targeted protein degradation may offer unique advantages over other modalities to study the transient/temporal changes in cellular signaling networks resulting from the acute depletion of proteins. Several CDK9 degraders have been reported over the last few years ^10–12^ that effectively degrade the kinase.

Here, we undertook our own efforts to design PROTAC molecules using the ultra-selective CDK9 inhibitor KI-ARv-03^13^, which was previously discovered in our group and the starting point for the clinical candidate KB-0742^14^ currently in Phase I/II clinical trials for patients with relapsed or refractory solid tumors or Non-Hodgkin Lymphoma (NCT04718675)^15–17^. We carried out transcriptional, proteomics, and phosphoproteomics profiling to comprehensively characterize the downstream effects and cellular adaptations that result from CDK9 degradation.

We report a selective and potent targeted CDK9 degrader that rapidly downregulates MYC levels. In addition, we explore cellular adaptations to CDK9 degradation and gain new insights into potential biomarkers indicative of contexts in which degradation might work better than inhibiting the kinase activity of CDK9. Our findings suggest that the selective degradation of CDK9 presents an attractive strategy for a robust attenuation of dysregulated transcription and could provide a therapeutic path for patients with aggressive and metastatic MYC-driven cancers.

## Results

### KI-CDK9d-32 is a potent CDK9 degrader with rapid kinetics

We designed and synthesized a first set of CDK9 PROTACs using KI-ARv-03 as the CDK9 recruitment moiety. The Small Molecule Microarray (SMM) platform used to discover KI-ARv-03 offers a suitable and intuitive starting point for the development of PROTAC molecules. Specifically, the original SMM screen involved compounds captured on isocyanate-coated glass slides, enabling rapid nomination of potential exit vectors for the attachment of linkers for PROTAC generation^18^. The intrinsic primary amine of the KI-ARv-03 molecule was used for linker connection. To construct the initial library of bivalent degraders, we used commercially available linkers to gain rough estimates of linker length and rigidity needed for degradation of CDK9 using KI-ARv-03 and Pomalidomide as recruiting element for the Cullin 4-Ring ubiquitin ligase cereblon (CRBN)^19,20^.

We evaluated the potential of these compounds to reduce CDK9 protein levels in MOLT-4 cells, a T-lymphoblast cell line where CDK9 is a known therapeutic target. The cells were treated with 1 µM of each compound for six hours. CDK9 protein levels were assessed using Western blot analysis.

Compound KI-CDK9d-08, a compound with a biphenyl linker (Figure 1A), stood out as the most effective PROTAC degrader from our preliminary library. We attributed this result to the rigidity of the biphenyl linker leading to favorable ternary complex formation with CDK9 and CRBN^23^.

**Figure 1:**
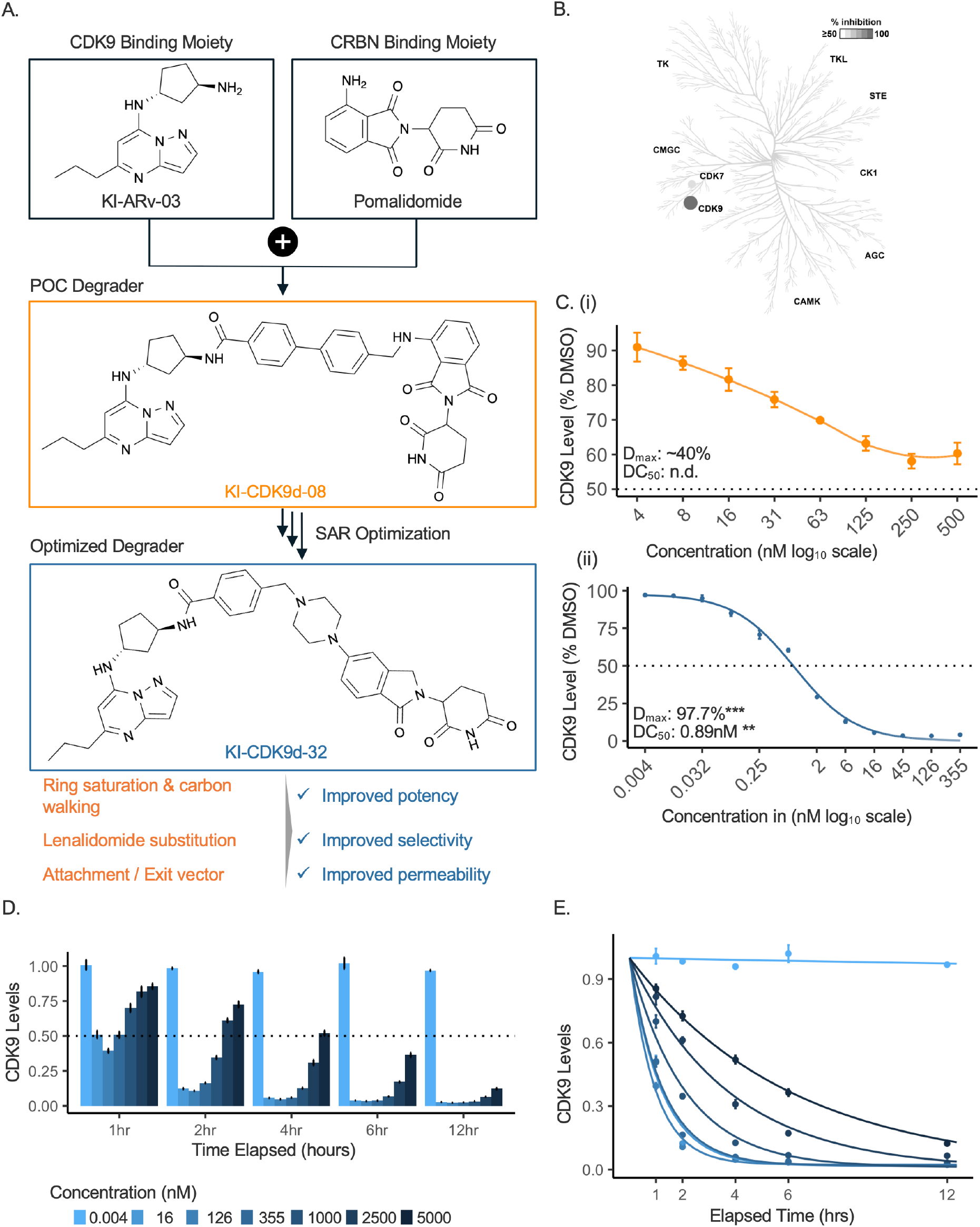
KI-CDK9d-32 is a potent CDK9 degrader with rapid kinetics. A) Summary of KI-CDK9d-32 development. KI-ARv-03 and pomalidomide were used as the CDK9 binding and CRBN binding moieties, respectively. Linker optimization followed current prevailing standards for achieving better properties. B) Kinase profiling of KI-ARv-03 (adapted from Richters et al.)^13^. C) Luminescence evaluation of endogenous HiBiT-tagged CDK9 levels in MOLT-4 cells after 4-hours of treatment with KI-CDK9d-08 (i) or KI-CDK9d-32 (ii). D) Kinetics evaluation of KI-CDK9d-32 at 1, 2, 4, 6 and 12 hours using luminescence readout of HiBiT-tagged CDK9 in MOLT-4. E. Kinetics profile of degrader KI-CDK9d-32. The data plotted are the same as D. The curves were fitted to the one-parameter exponential decay model^21^ and degradation rates at each concentration estimated. The kinetics profile follows that of a classical hook-effect^22^ model wherein there is a goldilocks effect, the maximal rate of degradation is bounded by a low and a high concentration. The rate at 16nM was 0.83Hr^-1^, the fastest rate of degradation (1.04Hr^-1^) for our experimental data occurs at 126nM, and drops to 0.80Hr^-1^ at 355nM when the hook-effect starts to dominate.

Considering the insights gained from our initial degrader library, we directed our subsequent optimization efforts toward the KI-CDK9d-08 scaffold (Figure 1A). We employed emerging design strategies from the targeted degradation space, such as linker optimization and IMiD substitutions, to enhance degradation efficiency and minimize unintended polypharmacology effects resulting from the neo-substrates targeted by CRBN^20^. This allows for a precise assessment of the biological effects driven primarily by CDK9 degradation. The field has seen significant advancements in the design and optimization principles for PROTAC molecules, especially after identifying a suitable starting point^20,24^. The incorporation of saturated rings, like piperidine and piperazine, into the linker has demonstrated improvements in potency and physicochemical characteristics, including solubility and metabolic stability^24^. Moreover, altering the attachment point of the linker to the phthalimide ring of the IMiD moiety and replacing pomalidomide with lenalidomide have proven to be effective modifications for diminishing the known off-target effects, mainly CRBN neo-substrates, of pomalidomide-based PROTACs^24^.

These strategies enabled us to successfully design a second-generation collection of degraders with enhanced properties from the KI-CDK9d-08 scaffold. We assessed the degradation efficiency of our optimized compounds in cell-based assays and selected KI-CDK9d-32 (D32), a sub-nanomolar degrader, as the leading candidate for generating biological insights. KI-CDK9d-32 achieves near 100% maximal degradation and a DC50 of 0.89nM following 4 hours of treatment in MOLT-4 cells (Figure 1C (ii)), making this, to our knowledge, the most potent CDK9 degrader reported. Consistent with the “hook-effect”, which is often observed with PROTACs and reflects the saturation of both protein members of the ternary complex with their respective binding elements and rendering the formation of a ternary complex unfavorable above a certain concentration threshold^25^, we observed limited degradation efficiency starting at 355 nM for the early time-points. Further assessment of the kinetics of degradation showed that this effect became increasingly negligeable as treatment time elapsed. CDK9 levels remained below 10% of the DMSO baseline for at least 12 hours after treatment (Figure 1D and E). We believed this convergence to complete CDK9 degradation over the course of our study to be the result of slower CDK9 resynthesis rates^22^.

### KI-CDK9d-32 is a highly selective and potent CDK9 degrader that induces rapid reduction of MYC protein levels

KI-CDK9d-32 exhibited a potent effect on the level of HiBiT-tagged CDK9. HiBiT is a CRISPR-mediated system that enables the development of luminescence-based assays to quickly assess the levels of endogenous proteins^26^. Given that selectivity is a critical consideration when targeting kinases, we then employed quantitative mass spectrometry to perform a comprehensive assessment of the selectivity and specific impacts of KI-CDK9d-32 on proteins in MOLT-4 cells. CDK9 and MYC exhibited the most pronounced reduction in protein levels. We observed an approximate 5-fold reduction in CDK9, which coincided with a 3-fold decrease in MYC protein levels (Figures 2A and 2B). Additionally, proteins such as AURKA, PLK1, and AURKB, which were found to be downregulated, are recognized for their role in a positive feedback loop with MYC^27,28^.

**Figure 2:**
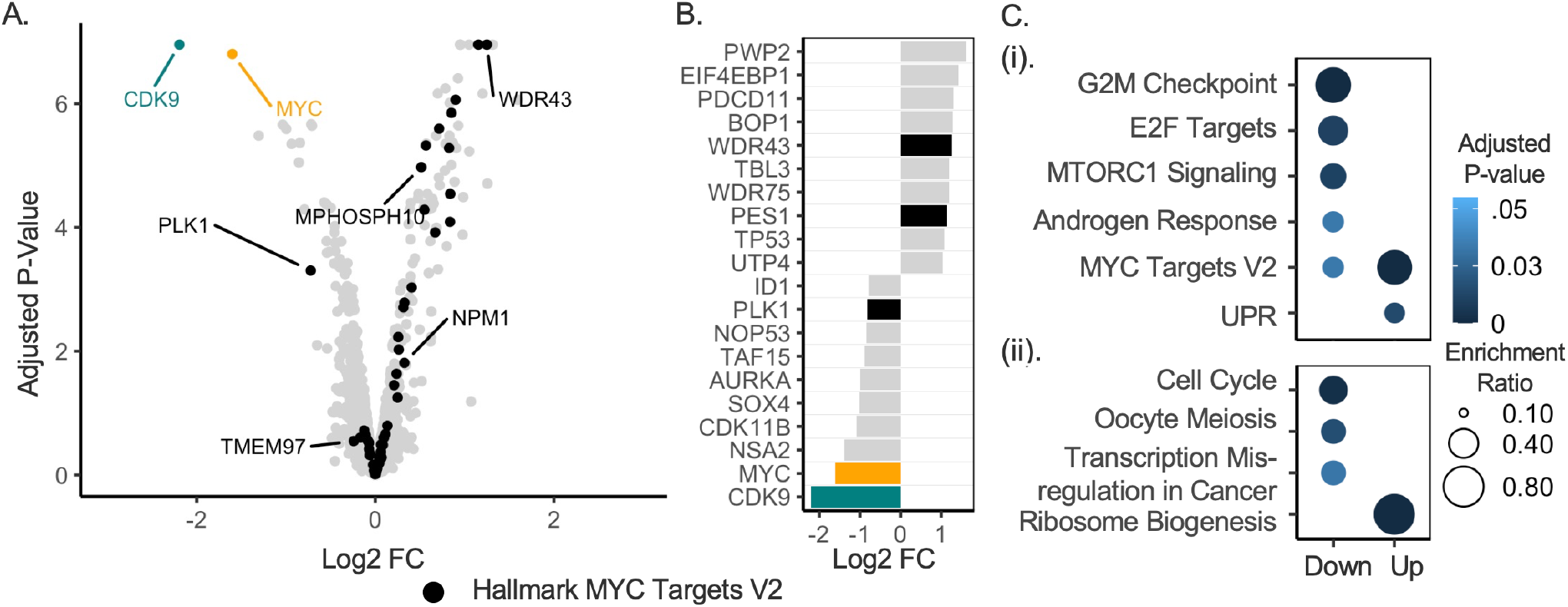
KI-CDK9d-32 is a highly selective and potent CDK9 degrader that induces rapid reduction of MYC protein levels. Quantitative mass-spectrometry assessment of the protein-level effects of KI-CDK9d-32. KI-CDK9d-32 has a robust effect on MYC-driven processes based on proteomics assessments in MOLT-4 cells. Cells were treated with DMSO or 50nM of KI-CDK9d-32 in four biological replicates. Protein lysates were harvested after 1 and 4 hours of exposure to the agents. A) Volcano plot representation of the 4 hour time-point. CDK9, MYC, and MYC target genes from one of the molecular signatures database (MSigDB) Hallmark collection are shown. B) Top 10 up and down regulated proteins from A. C) Enrichment analysis of the top 10% of genes that were differentially impacted – (i) enriched MSigDB Hallmark pathways and (ii) enriched KEGG pathways. The “Enrichment Ratio” is the ratio of observed significant proteins in a cluster belonging to a given pathway to the total number of proteins that could have been identified in that category in the background universe. In this case, we used all proteins identified in the experiment as the cell-specific background.

We conducted pathway over-representation analyses on the top 10% of proteins that were enriched (“Up”) or depleted (“Down”) following treatment. We performed these analyses using gene sets from the Molecular Signatures Database (MSigDB)^29^ and the Kyoto Encyclopedia of Genes and Genomes (KEGG)^30^ to identify pathways that were significantly enriched in the two clusters. The results demonstrated that proteins involved in cell cycle regulation, response to cellular stress and growth signals were depleted, with G2M checkpoint and E2F targets being the most significant negatively enriched pathways on treatment with the degrader, KI-CDK9d-32 (Figure 2C(i)). Depletion of proteins in these pathways indicates a strong inhibitory effect on cell cycle progression suggesting a coordinated cellular effort to halt proliferation and growth, in response to CDK9 degradation. Moreover, ribosome biogenesis is the most significantly KEGG enriched pathway for proteins in the “Up” cluster after the 4-hour treatment with 50nM KI-CDK9d-32 (Figure 2C(ii)). This positive enrichment of ribosomal proteins in the proteomic dataset was in alignment with nucleolar stress condition during which sequestration of nucleolar proteins to the cytoplasm occur, thus increasing their solubility^31^. This is further substantiated by the depletion of MYC and stabilization of TP53 observed following compound treatment (Figure 2A-B)^31^. These findings substantiate the degrader’‘s induction of a considerable disruption of the regulatory network associated with MYC.

### CDK9 degradation induces sustained disruption of MYC, and bypasses a known compensatory mechanism

Proteomics and gene set over-representation analyses suggested that degradation of CDK9 led to rapid and potent downregulation of MYC at the protein level, which is significant given MYC’‘s inherently fast turnover. This observation suggests extremely rapid CDK9 degradation kinetics that effectively shuts down MYC transcription. In contrast, previous studies have reported a compensatory increase in MYC levels following CDK9 pharmacological inhibition^2^. CDK9 degradation, on the contrary, seems to robustly bypass this resistance mechanism^32^ and as a result likely to have a pronounced impact on MYC-driven processes. To substantiate these results, we first performed qPCR assays to compare the effects of KI-CDK9d-32 and KB-0742 on the transcripts levels of CDK9 and MYC. As indicated previously by Lu et al.^2^, we observed an increase in MYC mRNA levels within 2 hours of CDK9 inhibition with 125nM and 1.2μM of inhibitor KB-0742. However, this paradoxical effect was not as pronounced at the 8-hour mark (Figure 3A).

**Figure 3:**
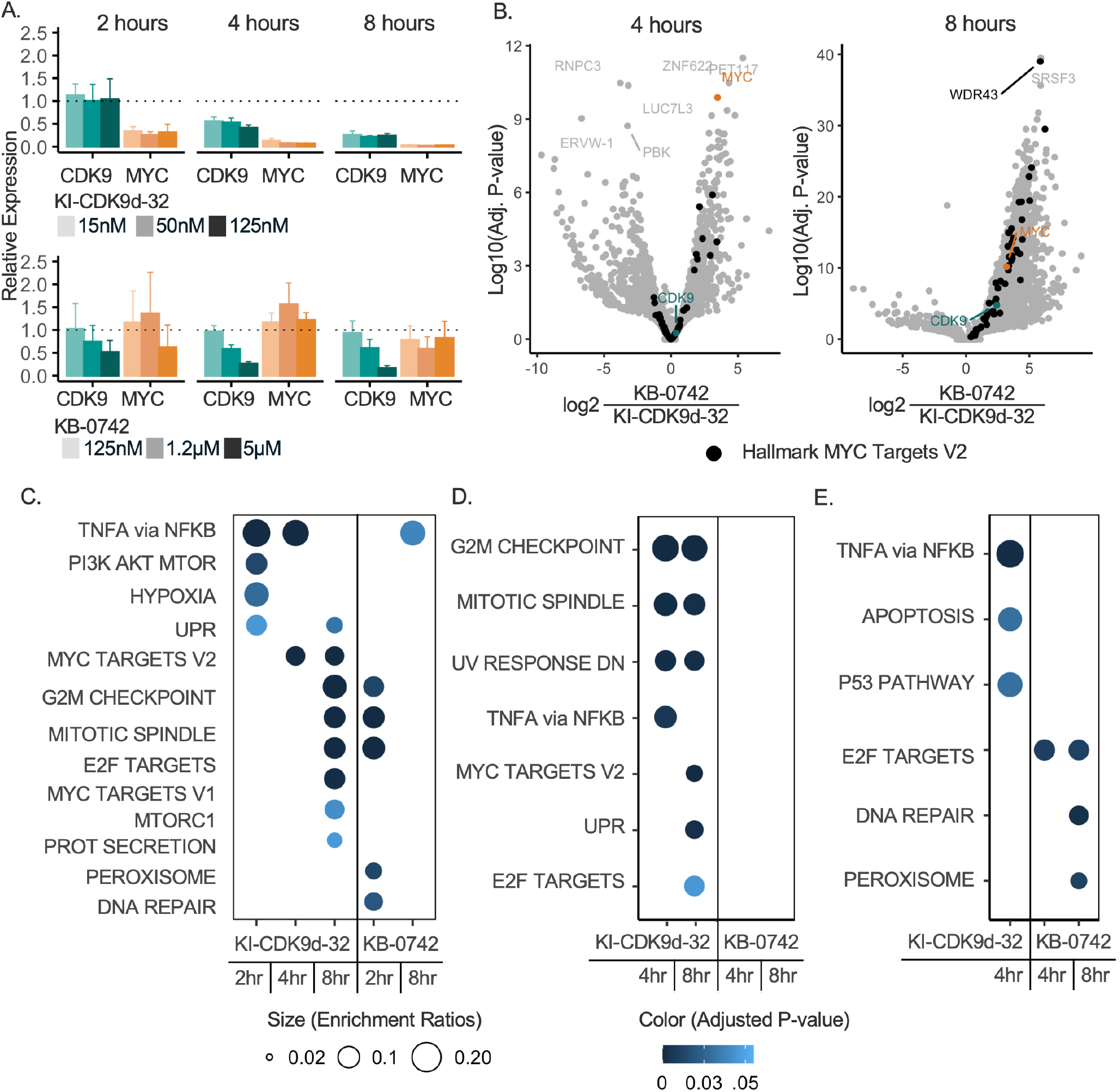
CDK9 degradation induces sustained disruption of MYC, and bypasses a known compensatory mechanism. *A)* RT-qPCR at the indicated time-points. Averaged across 3 biological replicates, and at least 3 technical replicates for all conditions. Degrader KI-CDK9d-32 repression of MYC mRNA levels is consistent in a time and dose-dependent manner (15nM – 125nM). Inhibition with KB-0742 shows a paradoxical increase in MYC transcripts at earlier time-points. The relative expression was determined using an average of the abundant POLR2A and 18SRNA as reference, and DMSO-treated samples as control. B) RNA sequencing evaluation of transcript levels following 4 and 8 hours of treatment with 1.2 μM of KB-0742 and 15nM of KI-CDK9d-32. LFC is the log2 of the ratio of Inhibitor/Degrader. The transcripts of genes with LFC > 0 are differentially downregulated by the degrader. C - E) Enrichment analysis of the top 20% of genes that were differentially downregulated in degrader versus inhibitor. As in B, LFC is the log2 of the ratio of Inhibitor/Degrader.

Treatment with KI-CDK9d-32 significantly suppressed transcription of MYC at all concentrations tested in both a dose- and time-dependent manner. The effect on CDK9 transcript levels did not show a similar drastic contrast between degradation and inhibition. Both treatments affected CDK9 transcript levels in a time and dose-dependent manner. However, the inhibitor shows this dose-dependence at the two highest concentrations (Figure 3A). This is consistent with the notion that high and persistent concentration of inhibitors is required for therapeutic efficacy^7^.

To identify and categorize other transcripts that display differential responses to the two pharmacological approaches, we carried out RNA sequencing experiments using synthetic RNA spike-ins to control for changes in the overall transcriptional state^33^. MOLT-4 cells were treated with 15nM and 1.2μM of KI-CDK9d-32 and KB-0742, respectively, for 2, 4, and 8 hours across four biological replicates. Consistent with our qPCR data, CDK9 degradation using 15nM KI-CDK9d-32 induces a rapid and sustained downregulation of MYC mRNA expression more than 6-fold relative to CDK9 inhibition (Figure 3B). Moreover, this reduction in MYC expression triggers a repression of CDK9 transcript levels in the RNA sequencing data, indicative of a potential positive feedback mechanism. This positive feedback mechanism becomes very apparent at the 8-hour mark, where CDK9 expression is reduced by about 5-fold by the degrader (Figure 3B, and Supplemental Figure S3.1).

To further analyze the transcriptional signature of each compound across the time-points tested, we performed over-representation enrichment analyses. Recognizing the limitations of relying solely on applying fixed fold-change thresholds, which may overlook biologically significant changes at earlier time-points, we employed a percentile-based significance filter. Specifically, included genes in the enrichment analyses were required to fall within the top 20% of absolute fold changes within a given condition tested. This allowed us to capture genes that show similar trends in their differential expression across timepoints even if their fold changes at earlier time-points are modest.

We observed subtle temporal differences between CDK9 degradation and inhibition. Degradation had a potent effect on early growth signaling and innate inflammation as evidenced by the negative enrichment of TNFA Signaling Via NFKB, and the Unfolded Protein Response pathways^34^ (Figure 3C). It appears that CDK9 degradation rapidly downregulated BRD4-mediated processes, which was expected given the structural and functional relationship between the P-TEFb and BRD4^34–36^. Moreover, the MYC Targets gene sets were differentially impacted. Comparing the effects of the two pharmacological approaches, we found that degradation had the strongest effect on the Hallmark “MYC Targets V2”, the set most correlated with ribosome biogenesis, with the tryptophan-aspartate repeat domain 43 (WDR43) one of the topmost downregulated. The effects of CDK9 Inhibition, on the other hand, was most pronounced against the DNA damage pathway (Figure 3B-C, Supplementary Figure S3.3).

In parallel to the above experiment in MOLT-4, we also evaluated the effects of KI-CDK9d-32 and KB-0742 treatment on transcriptional programs in two additional cell lines, the pancreatic adenocarcinoma cell line PSN-1 and the rhabdomyosarcoma line RH-4 to determine the presence of persistent signals across different cell lines. The experiments in these cell lines were carried out at the same concentrations as those described above, for durations of 4 and 8 hours.

The TNFA Signaling Via NFKB hallmark pathway was the most persistently differentially downregulated across all cell lines in the degrader treated samples relative to the samples treated with the inhibitor (Figure 3C – 3E). This pathway was consistently the earliest downregulated pathway in the samples treated with the degrader across all three cell lines. At the other end, the inhibitor appears most effective against genes involved in the maintenance of genomic stability. The DNA-repair pathway was significantly downregulated by the inhibitor in 2 of the three cell lines (Figure 3C-D, Supplementary Figure 3.3). These observations are reflective of important mechanistic differences driven in part by complete abrogation of CKD9 kinase and scaffolding functions by the degrader. The molecular consequence of which is a breakdown of the BRD4-P-TEFb complex, an insight that aligns with the rapid and sustained degradation of critical early response genes regulated by BRD4 (e.g., MYC and IFRD1) by the degrader^2,34,35^. Moreover, the kinetics of action of degradation versus inhibition of CDK9 is likely to be an important determinant. This underscores the significance of a probe with the rapid kinetics to enable a temporal assessment of the primary responses to inhibition versus degradation as observed in this study.

Overall, the degrader KI-CDK9d-32 appears to have more pronounced effects than the inhibitor KB-0742 on transcriptional repression of genes important for proliferation especially in MOLT-4 and PSN-1. This trend is reversed for most genes in the context of RH-4, where the inhibitor’‘s effects on MYC transcription appeared to be stronger. The subtle differences exhibited might be due to differences in transcriptional programs driven by lineage-specific core transcriptional regulatory circuitries^37,38^ (Supplementary Figures S3.4A – S3.4C).

### CDK9 degradation disrupts nucleolar homeostasis, a MYC-regulated process

Ribosome biogenesis, the process that governs protein synthesis in the nucleolus, is tightly regulated by MYC. The suppression of MYC network in cancer can lead to a collapse of ribosome biogenesis and induce widespread suppression of protein synthesis^39,40^. This can be especially disruptive to cancer cells given their elevated reliance on protein synthesis for aberrant proliferation. Numerous studies have demonstrated that CDK9 inhibition or silencing can have downstream effects on ribosomal biogenesis processes^10,41,42^. Consistent with these results, our proteomics and RNA sequencing analyses found ribosome biogenesis as the most significantly impacted cellular process following CDK9 degradation using KI-CDK9d-32. As seen with earlier studies^10^, we find the effect from CDK9 degradation on ribosome biogenesis to be stronger than that induced by inhibition (Figure 2C (ii), Supplementary Figure S3.1). Given the clear biological phenotype emerging from both protein-level and mRNA data, we carried out high resolution immunofluorescence microscopy to explore this further.

The Nucleophosmin protein (NPM1) and nucleolar RNA helicase 2 (DDX21) are key markers of nucleolar dynamics. NPM1 is a scaffold protein that plays a critical role in the assembly of the nucleolus and is localized at the nucleolar rim, the outer region of the nucleolus^43,44^. DDX21 is localized at the core of the granular compartment and is known to engage in several protein-protein interactions that drive ribosomal RNA metabolism^45^ (Figure 4B). Thus, we directly monitored the impact of the compounds on the nucleolar structural stability by staining HeLa cells for NPM1 and DDX21. Ribosome biogenesis has been extensively studied in HeLa cells, making them a well-suited model for these immunofluorescence experiments. KI-CDK9d-32 treatment destabilized nucleolar homeostasis as early as 2 hours, indicating that potent CDK9 degradation destabilizes nucleolar homeostasis by targeting, either directly or indirectly, the nucleolar rim, the outer layer of the nucleolus defined by NPM1 (Figure 4A). Interestingly, this phenotype seems to deviate from the previously described model of nucleolar stress. Unlike Actinomycin D, a known inducer of nucleolar stress, CDK9 degradation had little to no impact on the core granular compartment of the nucleolus. Possibly, CDK9 degradation is more potent toward the nucleolar pool of CDK9 and is highly disruptive to the interaction of RNA Polymerase II with ribosomal DNA^42,46^.

**Figure 4:**
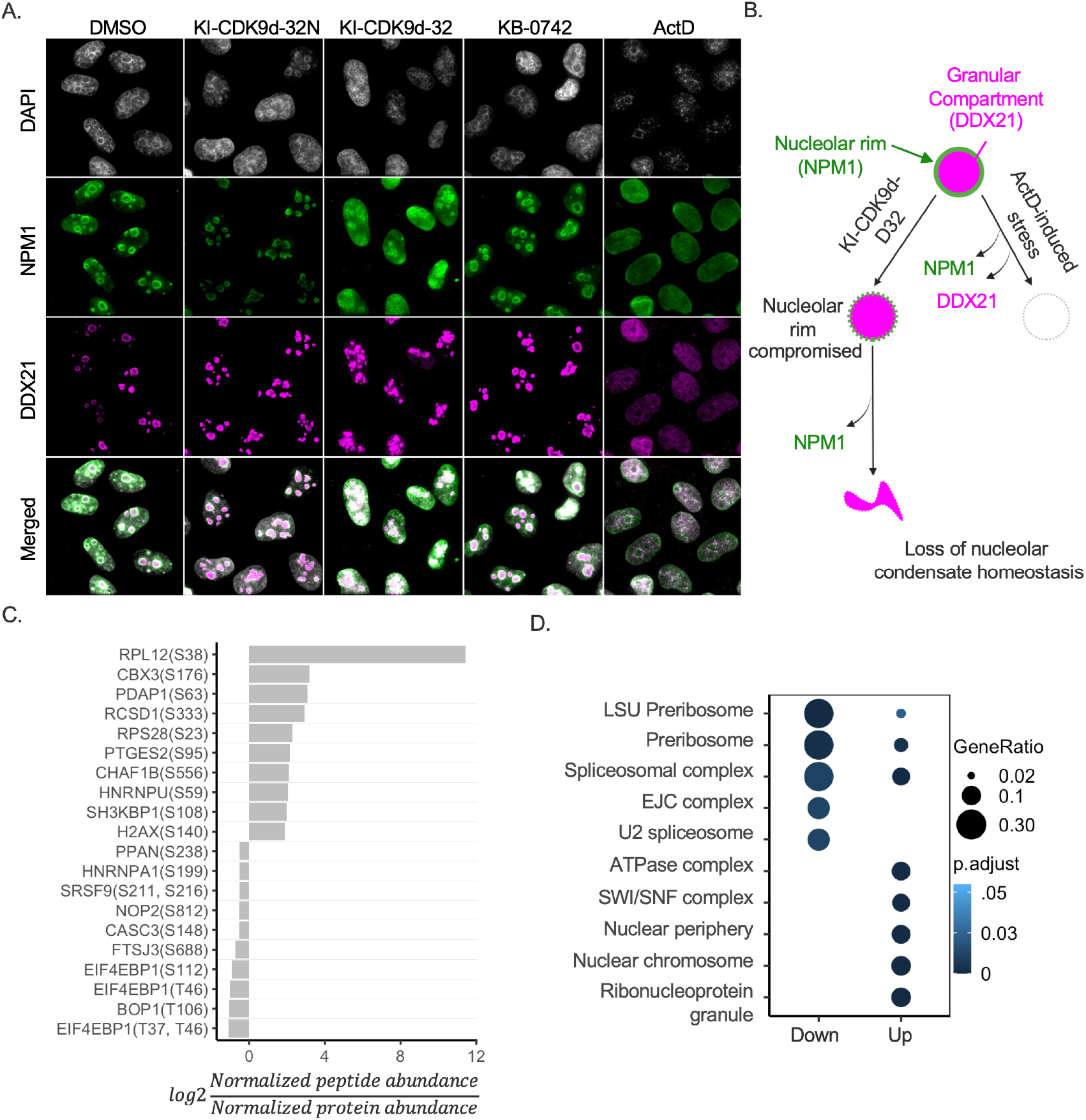
KI-CDK9d-32 potently destabilizes nucleolar homeostasis. A. Fluorescence imaging of HEK-293 cells following treatment with the inhibition, degrader, and relevant controls (DMSO, Negative Degrader, and Actinomycin D, a well-known inducer of nucleolar stress). Images were processed using ImageJ. KI-CDK9d-32N is the negative control of the degrader, lacking CRBN binding. Supplementary figure S4 provides additional details on the dose-response effects from both KB-0742 and KI-CDK9d-32. B. Illustration of the nucleolar compartments. C. Top 10 enriched and depleted phosphopeptides identified, using log2 fold change (log2FC) derived from the ratio of DMSO-normalized peptide to protein abundance in MOLT-4 cells treated with 50nM KI-CDK9d-32 for 4 hours. Analysis utilized four biological replicates and the Limma package for differential analysis post-median normalization. D. Gene over-representation analysis was conducted on the top 10% peptides showing log2FC > 0.5, focusing on the Gene Ontology cellular compartment.

Moreover, the phosphorylation state of nucleolar proteins plays an important role in the assembly of the nucleolus as well as its structural and functional integrity^47^. A recent study unveiled several phosphorylation sites that are crucial for the assembly of the nucleolus. Specifically, the phosphorylation of NPM1 at S254 and S260 was shown to limit its localization within the nucleolus^47^, a shift that is deleterious to the structural integrity of the nucleolus. Given these insights, we characterized the effects of CDK9 degradation at the phospho-proteome alongside our assessment of global differential changes in protein levels. We analyzed the abundance of phosphopeptides from the same four biological replicates as the quantitative proteomics experiment described earlier comparing KI-CDK9d-32 to DMSO-treated samples. The top differentially phosphorylated peptides were predominantly enriched in mRNA metabolism, ribosomal maturation, and protein synthesis processes (Figures 4C and 4D). For instance, we observe significant reduction in phosphorylated 4EBP1, the protein encoded by the EIF4EPB1 gene. This reduction in phosphorylation at specific residues, including Threonine-37/Threonine-46, is known to favor the interaction between 4EBP1 and eIF4E, a factor that is essential for the recruitment of the 40s ribosomal subunits to the 5’’ end of mRNA^48^. Thus, 4EBP1 binding to eIF4E induces an inhibitory effect on translation^28,48,49^.

Taken together, these results demonstrate that KI-CDK9d-32 mediated depletion of CDK9 and MYC levels leads to a robust disruption of MYC regulated processes. While MYC is known to regulate ribosome biogenesis at the transcriptional level, we show here that depleting the MYC transcriptional network causes the collapse of the nucleolus, the site of ribosome biogenesis.

### Cytotoxicity induced by KI-CDK9d-32 depends on CRBN and the activity of ABC transporters

We assessed the effect of KI-CDK9d-32 and its negative control KI-CDK9d-32N on the viability of an initial panel of three cell lines, MOLT-4, PSN-1, and RH-4. The viability effects were compared to KI-ARv-03, KB-0742, and a known CDK9 degrader, Thal-SNS-32. KI-CDK9d-32 induced the most pronounced cytotoxic effects relative to the inhibitors and degrader in the comparison set (Figure 5A and 5B).

**Figure 5:**
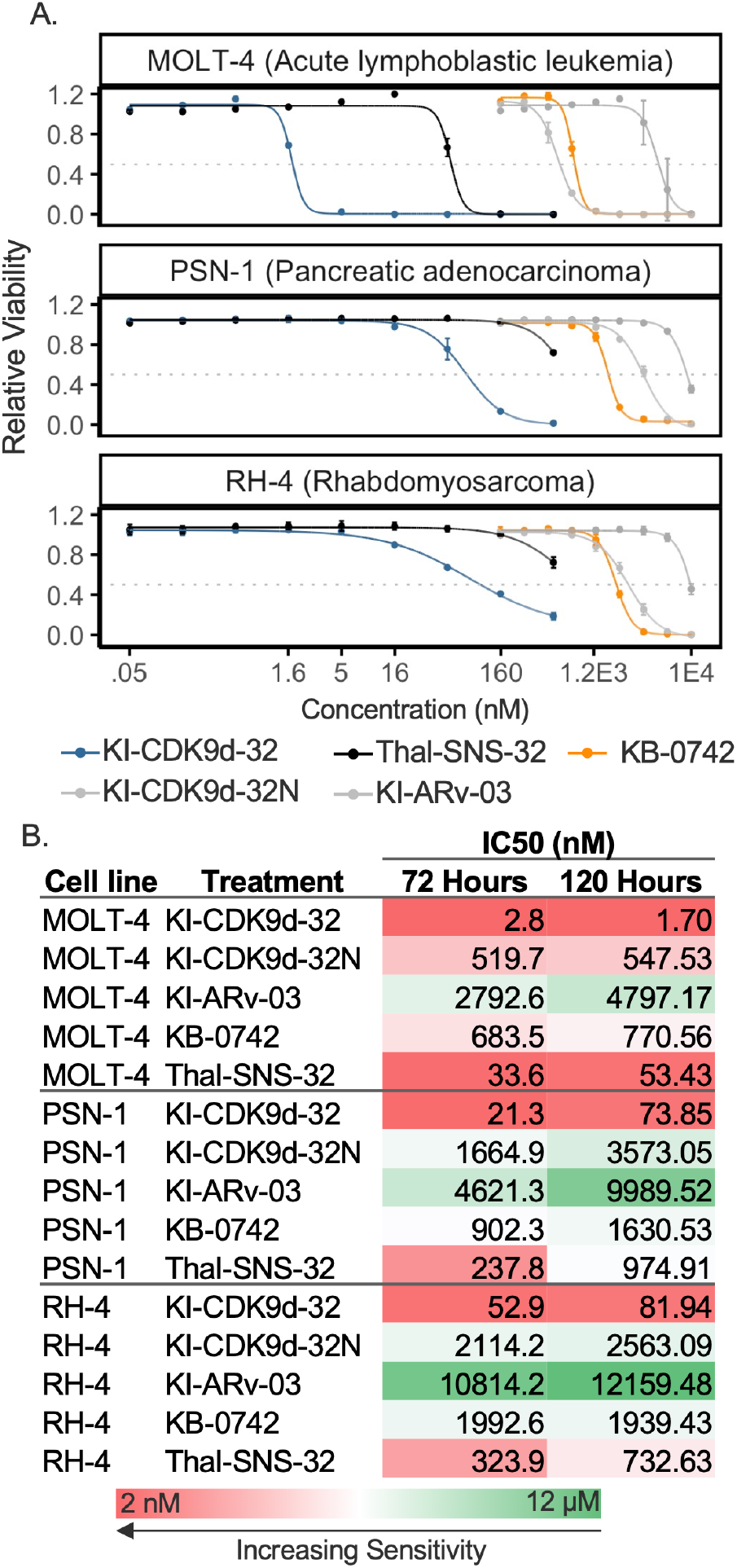
KI-CDK9d-32 demonstrates strong sensitivity in MOLT-4, PSN-1, and RH-4. (A) Dose-response curves representing cell viability of three cell lines (MOLT-4, PSN-1, and RH-4) measured 120 hours post-treatment with a panel of compounds: KI-CDK9d-32, KI-CDK9d-32N, KI-ARv-03, KB-0742, and Thal-SNS-32. X-axis is log transformed with concentration shown in anti-log for readability. Error bars indicate the mean ± standard deviation from n=3 technical replicates. (B) Tabulated IC50 values for the corresponding conditions at 72 and 120 hours post-treatment. Concentration ranges for degraders were 0– 500nM, while inhibitors and KI-CDK9d-32N were tested from 0–10μM. The dose response curves for the 72 hours endpoint as well as the structures of all compounds used are presented in accompanying Supplementary Figure S5.

As CDK9 perturbation continues to be pursued as an attractive therapeutic strategy for a variety of cancers, we were interested in determining which cellular models, if any, are likely to be most impacted by CDK9 degradation. We carried out broad cell line sensitivity profiling of ∼800 cancer cell lines through the Broad Institute’‘s PRISM sensitivity profiling platform^50^.

Given the pan-essential nature of CDK9^51^, we anticipated broad cytotoxic activity with both agents. Consistent with this assessment, treatment with KB-0742 and KI-CDK9d-32 had strong cytotoxic effects. The mean IC_50_ values were 926 nM and 82.6 nM for KB-0742 and KI-CDK9d-32, respectively. KB-0742 had a widespread cytotoxicity starting at 3.3 μM. The degrader exhibited a more selective cytotoxicity profile up to the maximal applied dose of 1.5μM (Figure 6A and supplementary Figure S6.1), about 100X the concentration used for the transcriptional profiling.

**Figure 6:**
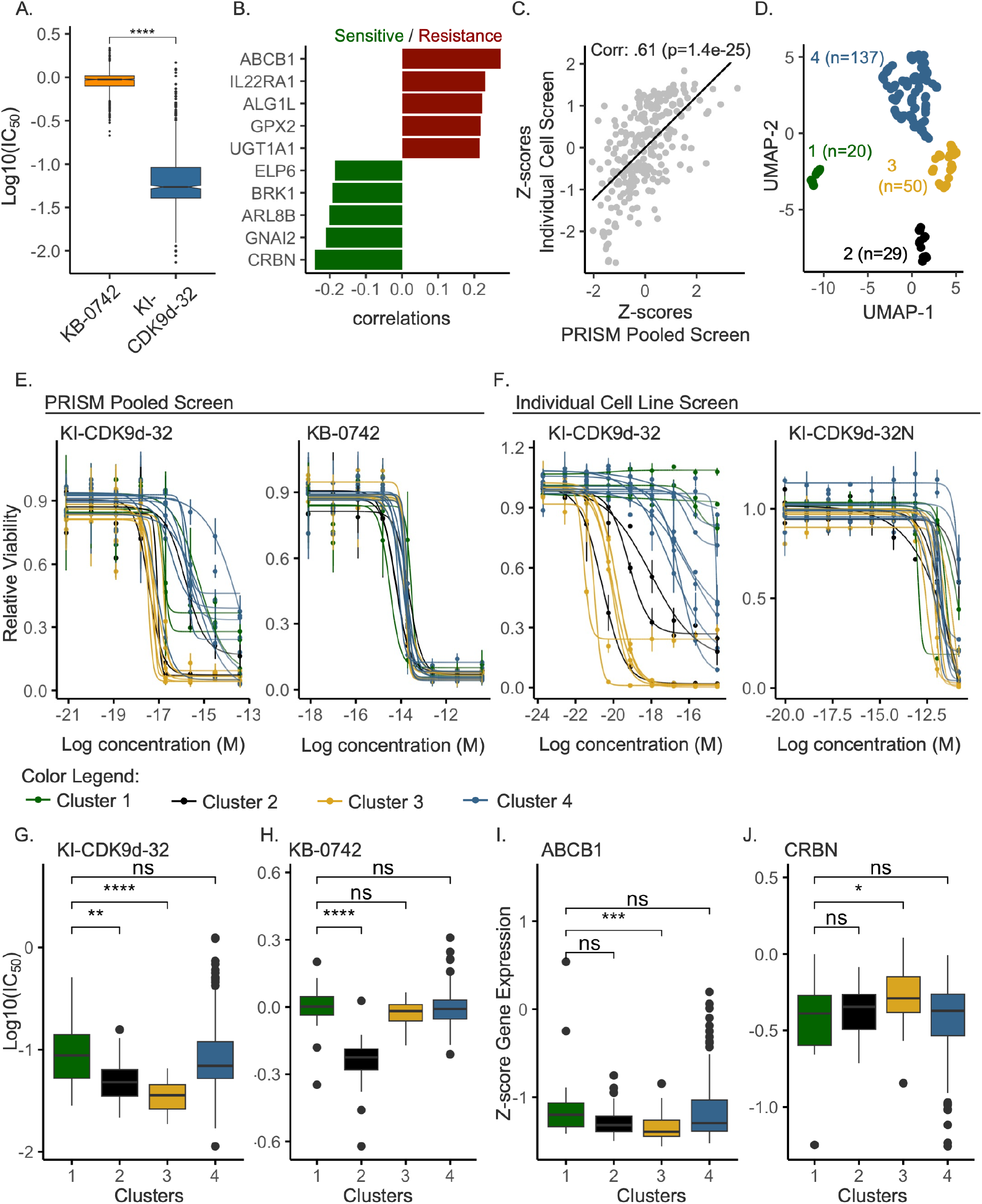
KI-CDK9d-32 has strong cytotoxic effects relative to the inhibitors used, but activity is limited in cells with high ABCB1 level. A) Distribution of the half-maximal inhibitory concentrations (IC50) resulting from a pooled screen of ∼800 cell lines through the Broad Institute’‘s PRISM platform. The PRISM platform offers a high throughput approach for compound screening in cancer cells derived from a variety of lineages (see supplement S6.2A for composition of lineages). KB-0742 and KI-CDK9d-32 were evaluated in 9-point three-fold dilutions series with top concentrations of 30µM and 1.5µM, respectively. B) Top significant gene expression biomarkers (q values < .05) of sensitivity to KI-CDK9d-32 (log2(IC50)). CRBN and ABCB1 were the top markers of sensitivity and resistance, respectively (supplement S6.2D provides an expanded figure). C) Correlation between the Z-scores of AUC values from the PRISM pooled screen and a secondary individual cell line screen. D) UMAP dimensionality reduction followed by HDBSCAN clustering of cell lines in both the PRISM pooled screen and the secondary individual cell line screen. The Z-score of IC50 and AUC values, relative metrics of compound sensitivity in the respective platforms, were used as inputs into UMAP to obtain a reduced dimensional data that used in HDBSCAN to obtain unbiased clustering of cell lines based on sensitivity patterns to the degrader and inhibitor. E, F).Dose-response plots of representative sets of cell lines from the clusters obtained in D. Viability values are relative to DMSO. Error bars represent the mean +/- sd from three and two replicates from the PRISM screen and secondary screen, respectively. The cell lines plotted in each cluster are listed in Supplementary Figure S6.2E. G,H.) Comparison of IC50 values of the full cluster constituents from the PRISM screen. I, J.) Show comparison to the expression profile of the strongest biomarkers from (B). For G through J., statistical comparisons were done using the pair-wise Wilcoxon rank-sum test between cluster 1 and the rest of the clusters, respectively. Statistical significance symbols as follows: ns: p > .05, *: p <= 0.05, **: p <= 0.01, ***: p <= 0.001, ****: p <= 0.0001

To determine the strongest biomarkers driving response to CDK9 degradation using KI-CDK9d-32, we examined the correlations between area-under the dose-response curve (AUC) values and the multi-omics features available through the DepMap portal following the standard PRISM analysis workflow^50^. CRBN expression, protein level, and copy number alteration consistently emerged as the most significant driver of sensitivity for degrader KI-CDK9d-32 (Figure 6B, Supplementary S6.2B). These observations are likely attributable to KI-CDK9d-32 as a lenalidomide-based degrader that induces the proximity of CDK9 and CRBN. Moreover, at the other end of the response spectrum, ABC-transporter mediated efflux activity surfaced as the top driver of resistance to KI-CDK9d-32: the gene expression and protein levels for ABCB1 were significantly correlated with higher AUC values (Figure 6B). This finding aligns with an earlier report on the role of ABCB1 in promoting resistance to PROTAC-based degraders^52^, and was consistent with results from a Madin-Darby canine kidney (MDCK) permeability assessment, a widely used assay in medicinal chemistry to determine whether compounds of interest are substrates for efflux transporters (Supplementary Table S1). In line with the proteomic and transcriptomic datasets discussed above, we also observed the ribosomal protein L15 (RPL15) as one of the top 10 biomarkers that correlated with response (low AUC values) to KI-CDK9d-32 (S6.2D).

To validate the above insights from the PRISM screen, we evaluated viability effects from the degrader against a set of 300 cell lines from a previously described cell repository at the MGH Cancer Center^53^. Cells were treated with either KI-CDK9d-32, or KI-CDK9d-32N, the negative control analogue of the degrader that differs only in the addition of methyl group to the glutarimide ring of lenalidomide and thereby preserving CDK9 binding while preventing recruitment CRBN. There was strong correlation between AUC values from the PRISM pooled and the non-pooled viability screening approaches (Figure 6C).

The impact of transporter activity complicated our efforts to sufficiently delineate cell lines differentially responsive to KI-CDK9d-32 and KB-0742 based on underlying mechanism. Despite this limitation of our data, we were able to identify clusters of cell lines that showed stronger sensitivity to one treatment over the other (Figure 6D,E,F). Supplementary Table S2 provides a full list of these cells and their response patterns. Interestingly, cells with high expression of ABCB1 were in general not responsive to both drugs (i.e., had the highest AUC values, Figure 6G-I).

## Discussion

Targeting MYC dependencies has been a significant therapeutic goal, promising to develop cures for various cancers. Approximately 15% of the global transcriptome is found to be dependent on the MYC regulatory network^54^. Dysregulation of MYC is estimated to occur in a vast number of cancers, with suggestions that up to 70% of cancers may exhibit MYC alterations^55^. The P-TEFb protein complex, which consists of CDK9 and CCNT1, plays a crucial role in the MYC network, affecting MYC’‘s transcription, activity, and stability^5,55–57^. As a result, numerous CDK9 inhibitors have entered clinical trials^6^.

The initial drug candidates, like flavopiridol, encountered significant challenges in clinical trials due to their broad targeting within the Cyclin-dependent Kinase protein family. The newer generation of CDK9 inhibitors, such as KB-0742, AZD4573, and VIP152, have demonstrated improved selectivity, and clinical trials are ongoing. However, the efficacy of CDK9 inhibition as a standalone treatment remains a subject of debate, particularly due to resistance mechanisms like the compensatory increase in MYC, or BCL-xL levels^2,58^. Combination regimens to address these limitations continue to be explored.

The emerging field of targeted protein degradation offers a novel avenue for probing CDK9 biology further and generating therapeutic hypotheses. Its appeal lies in the ability to inhibit both the enzymatic and scaffolding functions of target proteins, potentially preventing compensation or kinome rewiring^2,59^. Several groups have reported CDK9 degraders that exhibit greater selectivity than their precursor inhibitors. Here, we present the development a novel CDK9 degrader. We take this further by carrying out a comprehensive characterization of the effects of CDK9 degradation on transcription, global protein stability, and post-translation phosphorylation state. We found that CKD9 degradation induced a robust silencing of MYC transcription network. We see no signs of a compensatory increase in MYC mRNA expression over the study time points as previously documented with inhibition^2^. This suggests that CDK9 degradation could potentially offer additional therapeutic advantages over its inhibition.

Although certain cell lines with high transporter activity exhibited reduced sensitivity to KI-CDK9d-32, the overall sensitivity profile remained robust over inhibition. This is in line with the catalytic mechanism of action of degraders and underscores their potency at low intracellular doses. Further studies on KI-CDK9d-32 are necessary to confirm these findings in vivo. Additionally, the compound’‘s improved potency and physico-chemical properties highlight the potential for exploring novel delivery and targeting strategies. The findings presented here underscore the importance of continued exploration into the nuances of targeted degradation, not just as a means of drug development but as a fundamental lens through which we can better understand cellular regulation and disease progression.

## Acknowledgement

We thank Dr. Robert Wilson, Dr. Cameron Flower, and Dr. Tigist Tamir for their engagement in scientific and analytical discussions that strengthened the analyses and conclusions of the manuscript. This work was supported by the MIT Center for Precision Cancer Medicine (CPCM) and the National Science Foundation (Award #1845464). This work was supported in part by the Koch Institute Support (core) Grant 5P30-CA014051 from the National Cancer Institute. We thank the Koch Institute’‘s Robert A. Swanson (1969) Biotechnology Center for technical support, specifically the Barbara K. Ostrom (1978) Bioinformatics and Computing Core, the Genomics Core, and the Biopolymers and Proteomics Core Facilities. This work was also partially supported by a U54 Cancer Moonshot Grant (NCI-U54-CA231630). M.A.T. received funding support from a graduate fellowship from the Ludwig Center at MIT’‘s Koch Institute and the Alfred P. Sloan grant-funded University Center of Exemplarity Mentoring. M.A.T would like to thank Prof. Eric Fischer, and Prof. Bryan Bryson for relevant scientific discussions.

## Author Contributions

Conceptualization, A.N.K., A.R., M.A.T.; Medicinal Chemistry, K.M., A.R., J.U.; Investigation, M.A.T., K.M., Y.X., F.K.,S.S.L., R.S.; Bioinformatics & Visualization, M.A.T, C.W.; Data interpretation, M.A.T., A.N.K., E.C., F.W.; PRISM Screen, M.G.R. and J.A.R.; MGH Cancer Center Screen, C.J.O., A.K., R.I., K.G., A.K., S.O.; Writing – Original Draft, M.A.T; Writing – Review & Editing, all authors; Funding Acquisition & Supervision, A.N.K.

## Declaration of Interests

A.N.K is a scientific co-founder, scientific advisory board member, and equity holder in Kronos Bio. A.N.K, M.A.T., and K.M. have a patent application related to this work. K.M. and A.R. are current employees of Daiichi Sankyo and Plexium, respectively.

## Inclusion and Diversity

We support inclusive, diverse, and equitable conduct of research.

## RESOURCE AVAILABILITY

### Lead contact

Further information and requests for resources and reagents should be directly addressed to the lead contact, Angela N. Koehler.

### Materials availability

**KI-CDK9d-32** is available through http://koehlerlab.org/contact-us and upon completion of a material transfer agreement by the receiving institution.

### Data and code availability

- All data (RNA-Seq and Proteomics) will be deposited to the appropriate repositories prior to publication.
- This paper does not report original code.
- Any additional information required to reanalyze the data reported in this paper is available from the lead contact upon request.

## EXPERIMENTAL MODEL AND SUBJECT DETAILS

### Cell lines

Human acute lymphoblastic leukemia cells MOLT-4 (ATCC, CRL-1582), rhabdomyosarcoma cells RH-4 (Cellosaurus, CVL_5916), and pancreatic adenocarcinoma cells PSN-1 (ATCC, CRL-3211) were obtained from the Koch Institute’‘s Preclinical Modeling (ES Cell and Transgenics Facility) Core. Cell lines were maintained at 5% CO_2_ at 37°C. Cells were cultured in RPMI 1640 (Thermo Fisher Scientific, 11875093) + 10% fatal bovine serum (FBS). Cell lines were tested intermittently throughout studies using MycoAlert Mycoplasma Detection Kit (Lonza, Cat#LT07-418), generally a few days after thawing and immediately before significant studies (e.g. Proteomics, RNA-seq experiments).

## METHOD DETAILS

### Generation of HiBiT-tagged endogenous CDK9 in MOLT-4 cells

To generate MOLT-4 cells with an endogenous HiBiT-tagged CDK9, we utilized the CRISPR-Cas9 system to insert the HiBiT tag at the N-terminus of CDK9 immediately following the start codon^21,26^. In summary, Alt-R sgRNA and Cas9 Nuclease were combined to form a ribonucleoprotein complex (RNP), as per the guidelines provided by Integrated DNA Technologies (IDT). The transfection mixture was prepared by combining 5µL of the RNP complex, 2.4µL of 100µM Alt-R HDR donor oligonucleotides, 1.2µL of Alt-R Cas9 electroporation enhancer, 20µL of MOLT-4 cell suspension (containing 2x10^5 cells), and 1.4µL of PBS, reaching a total volume of 30µL per transfection. Electroporation was carried out using the 4D Nucleofector System (Lonza) and Nucleocuvette™ Strips, following the CA-137 program recommended by the manufacturer for MOLT-4 cells (Lonza, V4XC-2032). Post-electroporation, cells were incubated at room temperature for 5 minutes before being transferred to a six-well plate for cultivation. The electroporated cells were allowed a recovery period of one week. For the generation of monoclonal cell populations, cells were seeded in conditioned growth media at a limiting dilution to achieve approximately 0.5 cell per well in 96-well plates. After about two weeks of expansion, clones were screened for the insertion.

### CDK9 degradation assay

MOLT-4 cells expressing HiBiT-tagged CDK9 were cultured in RPMI 1640 medium (Thermo Fisher Scientific, 11875093) supplemented with 10% fetal bovine serum (FBS) and maintained in a 5% CO2, 37°C incubator. For degrader screening, dose-response, and kinetic studies, cells were seeded at a density of 10x10^3 cells per well in 96-well plates. The cells were treated with various compounds and a DMSO vehicle control in 100µL of growth medium. Following each treatment time point, replicate plates were equilibrated to room temperature, and an equal volume of 2X Nano-Glo HiBiT Lytic Reagent (Promega N3030)—comprising Nano-Glo HiBiT Lytic Buffer, Nano-Glo HiBiT Lytic Substrate, and LgBiT Protein—was added. The mixture was then agitated on an orbital shaker for 10 minutes to ensure thorough mixing, followed by a 10-minute incubation at room temperature to facilitate equilibration between LgBiT and HiBiT in the lysate. Luminescence was measured using the Tecan Infinite M200 plate reader.

### Immunofluorescence assay

Cells were seeded at 3x10^4^ cells per well in 24-well plates with glass coverslips (Fisher Scientific, 1254580) and incubated overnight for attachment. If applicable, drug treatments were administered for the specified durations. Cells were fixed with 4% paraformaldehyde directly in the media for 15 minutes at room temperature, followed by washes with phosphate-buffered saline (PBS) and stored at 4°C. Overnight blocking was performed at 4°C using PBSA buffer (PBS with 1% BSA, 0.1% Triton X-100, 0.05% sodium azide). Primary antibodies, DDX21 (Novus Biologicals, NB100-1718) and NPM1 (Abcam ab180607), were diluted to 1:200 in PBSA and applied to cells for 2 hours at room temperature. Secondary antibodies, Alexa Fluor 488 (Thermo Fisher Scientific, A-11008) and Alexa Fluor 647 (Thermo Fisher Scientific, A-27040), were diluted to 1:1,000 in PBSA and incubated for 1 hour at room temperature. Afterward, cells underwent three 5-minute washes with PBSA and two 5-minute washes with PBS, a brief water rinse, and were mounted on glass slides using ProLong Diamond antifade mountant (Invitrogen, P36961). Nuclei were stained with Hoechst 33342 (Thermo Fisher Scientific, 62249) at a 1:1000 dilution. Images were captured on a DeltaVision 2 TIRF microscope (60X objective) and processed accordingly.

### Quantitative Mass-spectrometry

#### Protein extraction

MOLT-4 cells were seeded in 6-well plates at a density of 2.4 × 10^6 cells per well, in 4 mL of RPMI-1640 supplemented with 10% FBS, approximately 12 hours prior to treatment. The cells were treated with either KI-CDK9d-32 or DMSO. A total of four biological replicates were prepared. At 1 and 4 hours post-treatment, the cells were lysed using RIPA buffer (Thermo Fisher Scientific, cat. 89900), which was supplemented with a cocktail of protease inhibitors (Thermo Fisher Scientific, cat. 87786), and phosphatase inhibitors (PhosSTOP, Millipore SIGMA, cat. 4906845001). The proteins in the lysates were precipitated using cold acetone (-20°C), following a standard protocol (Thermo Fisher Scientific TECH TIP #49), employing a single cycle of precipitation. The samples were then stored at -80°C until further analysis by mass spectrometry, as described below.

### Mass spectrometry sample preparation

#### Reduction, Alkylation, and Tryptic Digestion

Proteins were first reduced with 10 mM dithiothreitol (Sigma) for 1 hour at 56°C and then alkylated with 55 mM iodoacetamide (Sigma) for 1 hour at 25°C in the dark. The samples were diluted with 100 mM ammonium bicarbonate to decrease urea concentration to 1M. Proteins were digested overnight at 25°C with modified trypsin (Promega) using an enzyme-to-substrate ratio of 1:50 in 100 mM ammonium bicarbonate, pH 8.9. The reaction was stopped by adding formic acid (99.9%, Sigma) to achieve a final concentration of 5%. Samples were then desalted using Pierce Peptide Desalting Spin Columns (cat. # 89852).

#### TMT Labeling

Desalted samples underwent TMT labeling according to Thermo Fisher Scientific’‘s protocol. Samples were resuspended in 100 mM TEAB, vortexed, and briefly centrifuged. Anhydrous acetonitrile (20 µL) was added to the TMT label reagents, followed by vortexing, brief centrifugation, and a 5-minute dissolution period. Then, 20 µL of TMT reagents were added to each of the 100 µL samples, which were then vortexed and briefly centrifuged. The samples were incubated for 1 hour at room temperature. To quench the reaction, 5 µL of 5% hydroxylamine was added to each sample and incubated for 15 minutes. Equal volumes of each channel were combined and concentrated to dryness via speed-vac.

#### Phosphopeptide enrichment

A 10% aliquot of the combined TMT-labeled sample was reserved for HPLC fractionation to obtain the unmodified protein data. The remainder underwent phosphopeptide enrichment using the High-Select Fe-NTA Phosphopeptide Enrichment Kit (Thermo Fisher Scientific) as per the manufacturer’‘s instructions.

### Fractionation

#### HPLC fractionation of non-phosphopeptides

To acquire the unmodified protein data, 10% of the sample was fractionated using HPLC. The dried TMT-labeled peptides were resuspended in 10 mM TEAB and fractionated using an Agilent 1100 HPLC system with high pH buffers. Buffer A consisted of 10 mM triethylammonium bicarbonate (TEAB), and Buffer B was 10 mM TEAB in 99% acetonitrile. Fractions were collected every minute from the 10th to the 90th minute, resulting in 80 fractions. Every 15^th^ fraction was pooled resulting in 15 total fractions for further analysis. Specifically, the 15 fractions were dried, resuspended in 10 µL of 0.2% formic acid, and 10% of this mixture was injected into the LC-MS.

The peptides were resolved over the following gradient run:

| Time | % Buffer B | Flow rate µL |
| --- | --- | --- |
| 0 | 1 | 200 |
| 2 | 1 | 200 |
| 10 | 5 | 200 |
| 72 | 35 | 200 |
| 87 | 70 | 200 |
| 92 | 70 | 200 |
| 93 | 1 | 200 |
| 100 | 1 | 200 |

#### High pH fractionation of Phosphopeptides

Phosphopeptides were fractionated by Pierce High pH Reversed-Phase Peptide Fractionation Kit (Cat# 84868) per manufacturer’‘s instructions. Dried fractions were re-suspended in 3 μL of 0.2% formic acid, and 33% of this solution was injected into the LC-MS.

#### Liquid chromatography with tandem mass spectrometry measurement

The fractionated TMT-labeled tryptic peptides were separated by reverse-phase HPLC using a Thermo Fisher Scientific PepMap RSLC C18 column before nano-electrospray ionization in an Orbitrap Exploris 480 mass spectrometer (Thermo Fisher Scientific). The solvents used were 0.1% formic acid in water (Solvent A) and 0.1% formic acid in acetonitrile (Solvent B). The gradient ran from 1% Buffer B initially, increasing to 80% Buffer B, and then returning to 1% Buffer B over a total of 180 minutes. The mass spectrometer operated in data-dependent mode, targeting as many precursor ions as possible within a three-second cycle for MS/MS analysis.

### Protein identification and data pre-processing

#### Global quantitative mass spectrometry (Non-phosphopeptides)

The raw mass spectrometry data (.raw files) were analyzed using Sequest HT within Proteome Discoverer (Thermo Fisher Scientific). The search parameters included a 10 ppm mass tolerance for precursor ions and a 0.05 Da tolerance for fragment ion mass. Up to two missed cleavages by trypsin were allowed. Fixed modifications included carbamidomethylation of cysteine and TMT labeling of lysines and peptide N-termini. Variable modifications included oxidation of methionine, phosphorylation of serine, threonine, and tyrosine, loss of methionine at the protein N-terminus, acetylation of the protein N-terminus, and combined Met-loss plus acetylation at the N-terminus. Searches were conducted against both a human protein database (homo sapiens UniProt FASTA files) and an in-house contaminant database.

Peptide spectrum matches (PSMs) were exported from Proteome Discoverer (PD) and subjected to addition filtering and processing using a proteomic analysis workflow (PAW) described previously^60,61^. In summary, the PD exported PSMs were filtered based on the following default pipeline criteria: a q-value cutoff of 0.05, a mass deviation within ±20 ppm, and precursor isolation interference below 50%. PSMs were further filtered by charge state (2 to 4), peptide length (6 to 28 amino acids), and rank (considering only top-ranked hits). This filtered PSMs file was then used for high confident peptide and protein inference. At least 2 distinct peptides had to be present support a protein identification within a sample, and the minimum number of tryptic termini was 2, ensuring that peptides are fully tryptic. In a final processing step, proteins with insufficient evidence to be considered unique were grouped. Intensities are then added to the peptide and protein-level summaries and exported to R for quantitation and analysis.

#### Phosphopeptide

Phosphopeptide raw data were processed with Proteome Discoverer version 3.0 (Thermo Fisher Scientific), peptide and protein identification were performed with Mascot version 2.4 (Matrix Science), and phosphorylation site localization probabilities were assessed using ptmRS^62^ within Proteome Discoverer. Spectra were searched against the UniProt homo sapiens database. Search parameters were set to trypsin digestion with a maximum of two missed cleavages. Mass tolerance was set to 10 ppm at the MS1 level and 20 mmu at the MS2 level. Precursor ions and TMT reporter ions were removed from MS/MS spectra prior to searching using the non-fragment filter node in Proteome Discoverer. Cysteine carbamidomethylation, TMT-labeled lysine, and TMT-labeled peptide N-termini were set as static modifications; phosphorylation of tyrosine, serine, and threonine, and oxidation of methionine were set as dynamic modifications.

PSMs were exported from Proteome Discoverer and filtered to keep those with high quality: search engine rank of 1, delta mass within -10 to 10 ppm, expectation value <0.05, and ions score >15. PSMs missing values in more than 25% of TMT channels were discarded, remaining missing values were imputed with the minimum intensity observed in the dataset. Only peptides with an average abundance of 1000 or more across TMT channels, and with a ptmRS probability greater than 50 for at least one site were maintained. TMT reporter ion intensities were summed across PSMs sharing a common phosphopeptide sequence. To address the issue of redundancy, phosphopeptides derived from identical proteins with matching phosphorylation patterns were aggregated to streamline the dataset, ensuring each unique phosphosite was represented without unnecessary repetition. This aggregation was carried out by matching peptides based on their annotated sequences, modifications in master proteins, and positions in master proteins. Furthermore, motif analysis, leveraging the PhosphoSitePlus database^63^, was carried out to map phosphorylation sites to specific known motifs.

### Data / Differential analysis

#### Global quantitative mass spectrometry

The pre-processed global quantitative mass spectrometry data were loaded into R. For each unique protein, abundance values were summed. Normalization involved alignment of the datasets by applying a median-centering normalization across all samples. This process adjusted each sample’‘s abundance values by the ratio of the overall median to the sample’‘s median, thereby aligning median values across the dataset. Differential analysis was conducted using the Limma package in R, which facilitated the identification of differentially enriched or depleted proteins relative to the DMSO-treated control samples within each experimental condition. Gene set over-representation analysis was carried out on 10% of the most significant enriched or depleted proteins (adjusted p values < .05, and the absolute value of log 2 fold change > .5) to identify the cellular processes and functions most affected by the degrader. The clusterProfiler^64^ package from R was used. We used all proteins identified in the experiment as the universe for the over-representation analysis.

#### Phospho-proteomics

The pre-processed phosphopeptide data were loaded into R. For each unique peptide, abundance values were summed. Normalization involved two main steps: First, for drug-treated samples, abundance values were normalized against control samples within each experimental condition. Next, to align the datasets, a median-centering normalization was applied across all samples. This process adjusted each sample’‘s abundance values by the ratio of the overall median to the sample’‘s median, thereby aligning median values across the dataset. These normalized phosphopeptide data were then integrated with protein data, which underwent a similar normalization process, ensuring only peptides present in both datasets were retained. Differential analysis was conducted using the Limma package in R, which facilitated the identification of differentially phosphorylated peptides in relation to their overall protein levels. Gene over-representation analysis was carried out on the most significant enriched or depleted phosphopeptides (adjusted p values < .05, and the absolute value of log 2 fold change > .5) to identify the cellular compartments most impacted. The clusterprofiler package from R was used. The background universe used was all proteins identified in the global quantitative mass spectrometry.

### RNA extraction and RT-qPCR

MOLT-4 cells were cultured in RPMI-1640 medium supplemented with 10% FBS. Cells were seeded at a density of 2x10^6 cells/well in 6-well plates. Twelve hours after seeding, the cells in the assay wells were treated with KI-CDK9d-32 at concentrations of 15nM, 50nM, and 125nM; KB-0742 at concentrations of 125nM, 1.2μM, and 5μM; or DMSO (as a control) for durations of 2, 4, and 8 hours. At the end of the specified assay periods, RNA was extracted using the PureLink™ RNA Mini Kit (Invitrogen) and homogenized using the QIAshredder (Qiagen). Equal amounts of RNA were then used for cDNA synthesis with the iScript™ Advanced cDNA Synthesis Kit (Bio-Rad Laboratories, catalog number 1725038). RT-qPCR analysis was performed on the CFX384 Touch™ Real-Time PCR Detection System (Bio-Rad), following the manufacturer’‘s specified protocols and using SYBR Green (Roche/Kapa). The following primer pairs were used: MYC (5’‘-tcctcggattctctgctctc-3’‘/ 5’‘-tcttcctcatcttcttgttcctc-3’’), CDK9 (5’‘-ctgaagaaggtgctgatggaa-3’’ / 5’‘--agttgaccacattctcgtgtt-3’‘), 18s rRNA (5’‘-gctactggcaggatcaaccagg-3’’ / 5’‘-atgagccattcgcagtttcactg-3’‘), POLR2A (5’‘-ggatgatctggaatgctcag-3’’ / 5’‘-attccttgactccctccac-3’‘).

### RT-qPCR Data Analysis

The raw quantification cycle (Cq) values from RT-qPCR were exported from the CFX384 Real-Time PCR Detection System and imported into R software for statistical analysis. The median Cq values were computed from a minimum of three technical replicates for each unique combination of treatment, time point, specific gene, and concentration. To normalize gene expression, the delta Cq (ΔCq) for each condition was calculated by subtracting the Cq value of the POLR2A and 18S rRNA reference gene from the Cq value of the target gene. The Cq values of the two references were averaged per condition and this value used as the reference. The delta delta Cq (ΔΔCq) was then determined by subtracting the ΔCq of the DMSO control (the normalization control) from the ΔCq of each experimental condition. This step quantifies relative gene expression levels. The fold change in expression for each condition was defined as 2^(-ΔΔCq). The average fold change in expression from three biological replicates was used to compare the effects of the compounds on gene expression.

### RNA-seq

#### RNA extraction, synthetic RNA spike-in

Four biological replicates were used for these experiments. For each biological replicate, 10^3^ cells were seeded and treated with treated with KB-0742, KI-CDK9d-32, or DMSO. At the appropriate endpoint (2, 4, or 8 hours after treatment), cells were immediately lysed in lysis buffer (from PerkinElmer’‘s chemagic RNA Tissue 360 H96, VD200615) supplemented with BME flash frozen and stored at -80c. After all biological replicates were collected, samples were thawed in deep 96-well plates. An equal amount (∼30pg) of Lexogen’‘s ERCC spike-in mix (SIRV-3) were added at this point following manufacturer’‘s recommendations. The PerkinElmer’‘s Chemagic 360 was used for bead-based RNA extraction following manufacturer’‘s standard protocol.

RNA was quantified using an Agilent Fragment Analyzer, purified using 2X RNA SPRI beads, and 1ng of total RNA was used for Illumina library preparation. Libraries were prepared using a reduced volume version of SMART-seq2^65^ performed on an SPT Mosquito HV using 10 cycles of amplification for cDNA generation and 16 cycles for library generation using NexteraXT (Illumina). Libraries were sequenced on an Illumina NextSeq500 using paired end 38nt reads.

### RNA-seq processing and differential analysis

RNA-seq data was used to quantify transcripts from the hg38 human assembly with the gencode version 43 annotation using the nf-core/rnaseq workflow revision 3.12.0^66^. The genomic target included the LexogenSIRVData spike-in reference. Gene level summaries were prepared from the star_salmon quantitation using tximport version 1.28.0^67^ running under R version 4.3.0 (R Core Team 2021) with tidyverse version 2.0.0^68^. Differential expression analysis was done with DESeq2 version 1.40.1^69,70^ using apeglm log fold change shrinkage^71^. The R image used for this work is available (docker://bumproo/rnaseqclass23).

For each of the three cell lines, MOLT-4, PSN-1, and RH-4, protein coding genes and genes with sufficient variance and expressions across all conditions (variance > 0.1, and mean > 0.1) were used for downstream analyses. Enrichment analysis was conducted on the top 20% of genes that were differentially expressed when comparing the degrader directly with the inhibitor. The clusterprofiler R package^64^ was used for the over-representation analysis against the Molecular Signature Database Hallmark gene set. All protein coding genes in each dataset was used as background for that specific cell line to preserve any cell-specific expression. Preranked Gene Set Enrichment Analysis^72^ was done using javaGSEA version 4.3.2 with msigDb v2023.2 human gene sets^73^.The ranking metric used for GSEA was the Wald statistic produced by DESeq2 from differential expression tests run using data normalized to SIRV spike-ins.

### Cell viability assessment of MOLT-4, RH-4, and PSN-1 cells

After 12 hours post-seeding, cells in assay wells were treated with vehicle (DMSO) or compound stocks dissolved in DMSO. The cells were seeded at the following densities to accommodate for growth over the experimental period: MOLT-4 at 6.5x10^3^ and 3.0x10^3^, PSN-1 at 2.0x10^3^ and 1.0x10^3^, and RH-4 at 3x10^3^ and 1.5x10^3^ cells per well for 72-hour and 120-hour endpoint readings, respectively. Treatments included degraders (e.g., KI-CDK9d-32, Thal-SNS-32) at concentrations up to 500nM, inhibitors (e.g., KI-CDK9d-32N, KI-ARv-03, and KB-0742) up to 10µM, and the DMSO control. The percentage of DMSO by volume was kept below 0.4%. Cell viability was determined using the CellTiter-Glo assay (Promega), with luminescence measured by a Tecan Infinite M200 plate reader at the specified time points. Viability was determined as the normalized luminescence to the DMSO control for each cell line and replicate (n=3 replicates). The dose-response curves were fitted to a four-parameter log-logistic model to estimate IC50 values for both the 72-hour and 120-hour endpoints using the drc package in R.

### High throughput cellular sensitivity profiling

#### PRISM Pooled Cell Sensitivity Screen

The PRISM cell sensitivity screen was carried out as previously described^50^. In summary, approximately 800 barcoded cell lines in pools of 20-25 were thawed and plated into 384-well plates (1250 cells/well for adherent cells, 2000 cells/well for suspension or mixed suspension/adherent pools). Cells were treated in triplicate with threefold dilutions starting at 30 μM and 1.5 μM of KB-0742 and KI-CDK9d-32, respectively. Cells were incubated for 120 hours, then lysed. Each cell’‘s barcode was read out by mRNA based Luminex detection as described previously and input to a standardized R pipeline (https://github.com/broadinstitute/prism_data_processing) to generate viability estimates relative to vehicle treatment and fit dose-response curves. The IC_50_ and area under the dose-response-curve (AUC) values were used as metrics of therapeutic potency in cell lines and correlated with gene expression (transcripts/million) and proteomics data as annotated in the Cancer Cell Line Encyclopedia (CCLE).

#### MGH Cancer Center Individual Cell Line Sensitivity Screen

Cell lines are maintained as part of the MGH Center for Molecular Therapeutics for drug sensitivity profiling^53^ (Garnett et al, Nature 2012). All lines were tested for mycoplasma and grown in either RPMI or DMEM/F12 media with 10% fetal bovine serum. Cells were seeded into 384-well plates one day prior to drug addition. Cells were treated with vehicle (DMSO) or compound stocks dissolved in DMSO with a PerkinElmer JANUS workstation. Following drug addition, cells were incubated for 5 days. Cell viability was then determined using the CellTiter-Glo assay (Promega), with luminescence measured by a PerkinElmer Envision plate reader. Viability was determined as the normalized luminescence to DMSO control for each cell line (n=2 replicates). Dose-response curves were fitted to a four-parameter log-logistic model using SciPy scipy.optimize.curve_fit to estimate IC_50_ and AUC values.

#### Sensitivity Analysis and Clustering

To generate cell sensitivity clusters for KI-CDK9d-32 and KB-0742, AUC and IC50 values for cells in both the pooled and Individual cell line screens were independently standardized using z-scores (i.e., by scaling the mean to 0 and the standard deviation to 1). This led to relative AUC and IC50 values, controlling for systematic differences attributable to varying dose ranges. The scaled values were subjected to UMAP-based dimensionality reduction followed by hdbscan clustering. The UMAP parameters used were n_neighbors = 5, min_dist = .01, metric = Euclidean. For hdbscan, the default minPts=10 was used.

## Chemical synthesis and characterization

### 4’‘-(((2-(2,6-dioxopiperidin-3-yl)-1,3-dioxoisoindolin-4-yl)amino)methyl)-N-((1R,3R)-3-((5-propylpyrazolo[1,5-a]pyrimidin-7-yl)amino)cyclopentyl)-[1,1’‘-biphenyl]-4-carboxamide (KI-CDK9d-08)

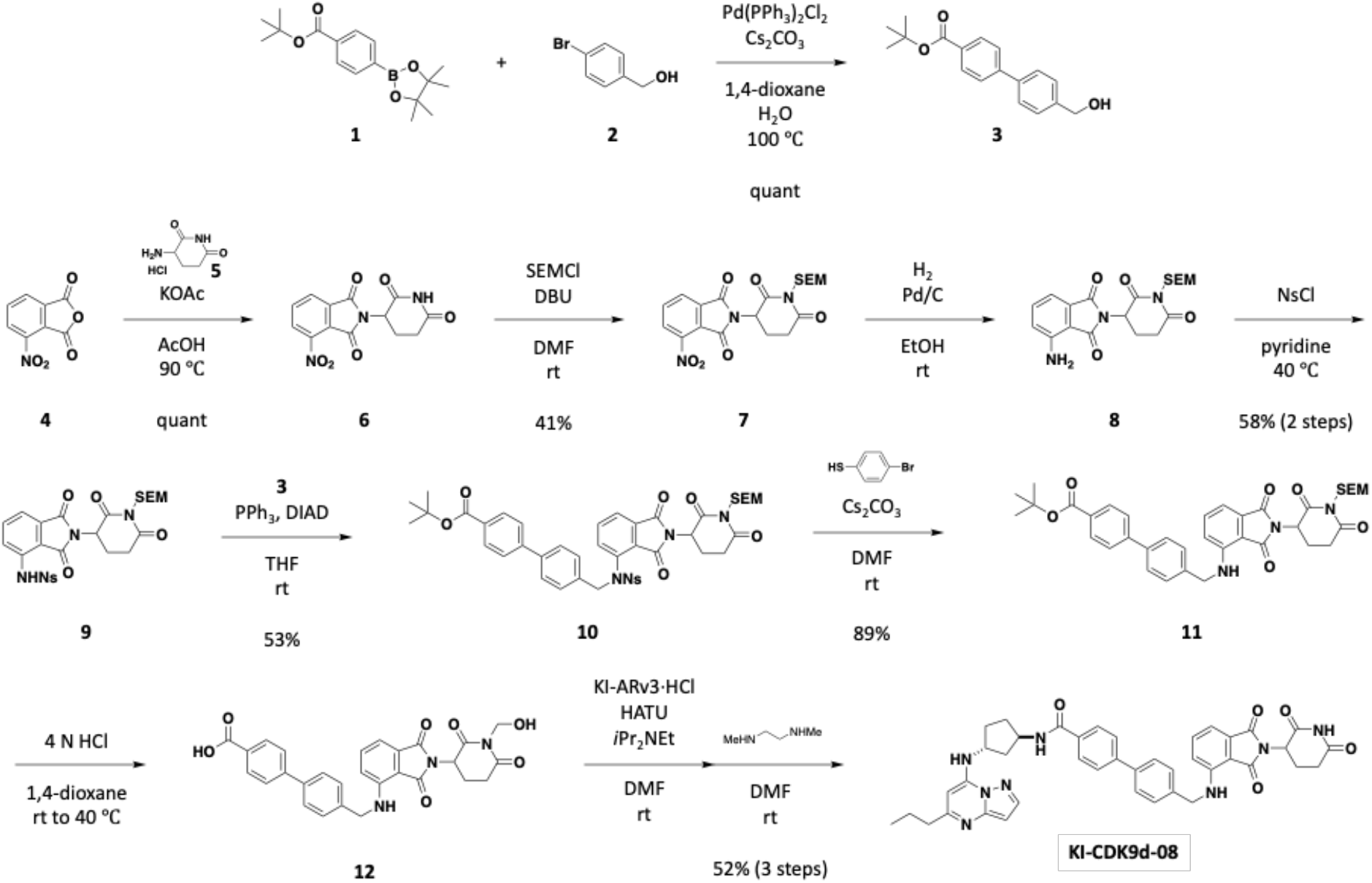

#### *tert*-butyl 4’‘-(hydroxymethyl)-[1,1’‘-biphenyl]-4-carboxylate (3)

A mixture of 4-(*tert*-Butoxycarbonyl)phenylboronic acid, pinacol ester (**1**, 1.00 g, 3.29 mmol), (4-Bromophenyl)methanol (**2**, 615 mg, 3.29 mmol), Pd(PPh_3_)_2_Cl_2_ (115 mg, 0.164 mmol), and Cs_2_CO_3_ (2.14 g, 6.57 mmol) in 1,4-dioxane (8.00 mL) and H_2_O (2.00 mL) was stirred at 100 °C. After 3.5 hours, the mixture was filtered through a pad of Celite and rinsed with ethyl acetate. The filtrate was concentrated to give a crude material, which was purified by silica gel column chromatography (hexane:ethyl acetate = 90:10 to 60:40) to yield the title compound as a white solid (997 mg, quant). ^1^H NMR (500 MHz, CDCl_3_) δ 8.05 (d, *J* = 8.5 Hz, 2H), 7.62 (dd, *J* = 8.3, 6.6 Hz, 4H), 7.46 (d, *J* = 8.0 Hz, 2H), 4.76 (s, 2H), 1.62 (s, 9H). ^13^C NMR (126 MHz, CDCl_3_) δ 165.82, 144.87, 140.89, 139.68, 130.96, 130.09, 127.65, 127.58, 126.95, 81.21, 65.13, 28.37. QToF HRMS m/z: calcd for C18H20NaO3^+^ [M+Na^+^] = 307.1305; Found 307.1311.

#### 2-(2,6-dioxopiperidin-3-yl)-4-nitroisoindoline-1,3-dione (6)

A mixture of 4-nitroisobenzofuran-1,3-dione (**4**, 2.00 g, 10.4 mmol) and 3-aminopiperidine-2,6-dione hydrochloride (**5**, 1.88 g, 11.4 mmol) and KOAc (3.15 g, 32.1 mmol) in AcOH (20.8 mL) was stirred at 90 °C overnight. The mixture was concentrated, and the resulting solid material was washed with methanol. The title compound was obtained as a gray solid (3.32 g, quant). ^1^H NMR (500 MHz, DMSO) δ 11.17 (s, 1H), 8.35 (d, *J* = 8.0 Hz, 1H), 8.24 (d, *J* = 7.6 Hz, 1H), 8.12 (t, *J* = 7.8 Hz, 1H), 5.20 (dd, *J* = 12.9, 5.3 Hz, 1H), 2.89 (ddd, *J* = 17.2, 13.9, 5.4 Hz, 1H), 2.66 – 2.57 (m, 1H), 2.56 – 2.45 (m, 1H), 2.12 – 2.03 (m, 1H). ^13^C NMR (126 MHz, DMSO) δ 172.74, 169.52, 165.19, 162.54, 144.44, 136.84, 133.02, 128.89, 127.32, 122.57, 49.44, 30.88, 21.74. QToF HRMS m/z: calcd for C13H9KN3O6^+^ [M+K^+^] = 342.0123; Found 342.0127.

#### 2-(2,6-dioxo-1-((2-(trimethylsilyl)ethoxy)methyl)piperidin-3-yl)-4-nitroisoindoline-1,3-dione (7)

DBU (2.22 mL, 14.9 mmol) and SEMCl (1.98 mL, 11.2 mmol) were added to a stirred solution of **6** (2.26 g, 7.45 mmol) in DMF (24.8 mL) at room temperature. After 2 hours, the mixture was quenched by adding saturated aq. NH_4_Cl, and extracted with ethyl acetate. The combined organic phase was dried over anhydrous Na_2_SO_4_. Filtration and concentration gave the crude material, which was purified by silica gel column chromatography (hexane:ethyl acetate = 90:10 to 50:50) to yield the title compound as a white solid (1.31 g, 41%). ^1^H NMR (500 MHz, DMSO) δ 8.36 (d, *J* = 8.1 Hz, 1H), 8.24 (d, *J* = 7.4 Hz, 1H), 8.13 (t, *J* = 7.8 Hz, 1H), 5.34 (dd, *J* = 13.1, 5.4 Hz, 1H), 5.08 (s, 2H), 3.52 (dtd, *J* = 30.6, 9.7, 6.4 Hz, 2H), 3.03 (ddd, *J* = 17.3, 14.0, 5.4 Hz, 1H), 2.85 – 2.77 (m, 1H), 2.60 – 2.51 (m, 1H), 2.15 – 2.07 (m, 1H), 0.91 – 0.77 (m, 2H), -0.02 (s, 9H). ^13^C NMR (126 MHz, DMSO) δ 171.56, 169.47, 165.11, 162.45, 144.46, 136.86, 132.98, 128.91, 127.30, 122.53, 68.35, 65.99, 49.98, 31.09, 20.73, 17.46, -1.38. QToF HRMS m/z: calcd for C19H27N4O7Si^+^ [M+NH_4_^+^] = 451.1644; Found 451.1648.

#### *N*-(2-(2,6-dioxo-1-((2-(trimethylsilyl)ethoxy)methyl)piperidin-3-yl)-1,3-dioxoisoindolin-4-yl)-2-nitrobenzenesulfonamide (9)

A mixture of **7** (660 mg, 1.52 mmol) and Pd/C (10%, 81.0 mg, 0.076 mmol) in ethanol (7.60 mL) was stirred under hydrogen atmosphere at room temperature overnight. The mixture was filtered through a pad of Celite, then concentrated. The resulting crude material was used in the next step without further purification. To a stirred solution of the crude material in pyridine (5.10 mL), 2-nitrobenzenesulfonyl chloride (1.01 g, 4.58 mmol) was added, then the mixture was stirred at 40 °C overnight. After cooling to room temperature, the mixture was quenched by adding 10 drops of H_2_O. Concentration and purification by silica gel column chromatography (hexane:ethyl acetate = 80:20 to 40:60 including 1% Et_3_N, then 100% ethyl acetate) gave the title compound as a yellow solid (521 mg, 58%, 2 steps). ^1^H NMR (500 MHz, CDCl_3_) δ 9.66 (d, *J* = 4.1 Hz, 1H), 8.19 (dt, *J* = 7.3, 1.8 Hz, 1H), 8.09 (dd, *J* = 8.6, 3.5 Hz, 1H), 7.92 (dt, *J* = 7.7, 1.9 Hz, 1H), 7.80 – 7.72 (m, 2H), 7.70 (t, *J* = 7.9 Hz, 1H), 7.53 (dd, *J* = 7.3, 2.2 Hz, 1H), 5.24 (s, 2H), 4.99 – 4.91 (m, 1H), 3.67 – 3.53 (m, 2H), 3.05 – 2.92 (m, 1H), 2.85 – 2.72 (m, 2H), 2.16 – 2.05 (m, 1H), 0.98 – 0.88 (m, 2H), -0.01 (s, 9H). ^13^C NMR (126 MHz, CDCl_3_) δ 170.77, 168.64, 168.08, 166.43, 148.18, 136.54, 135.93, 134.86, 133.08, 132.78, 132.23, 131.14, 126.15, 122.68, 119.13, 116.86, 69.38, 67.59, 50.22, 32.08, 21.84, 18.17, -1.32. AccuTOF DART HRMS m/z: calcd for C25H32N5O9SiS [M+NH_4_^+^] = 606.1685; Found 606.1721.

#### *tert*-butyl 4’‘-(((*N*-(2-(2,6-dioxo-1-((2-(trimethylsilyl)ethoxy)methyl)piperidin-3-yl)-1,3-dioxoisoindolin-4-yl)-2-nitrophenyl)sulfonamido)methyl)-[1,1’‘-biphenyl]-4-carboxylate (10)

Triphenylphosphine (80.2 mg, 0.306 mmol) and diisopropyl azodicarboxylate (60.2 mL, 0.306 mmol) were added to a stirred solution of **9** (150 mg, 0.255 mmol) and **3** (87.0 mg, 0.306 mmol) in THF (2.55 mL) at room temperature. After 22 hours, the mixture was quenched with H_2_O, and extracted with ethyl acetate. The combined organic layer was dried over anhydrous Na_2_SO_4_. Filtration and concentration gave a crude material, which was purified by silica gel column chromatography (hexane:ethyl acetate = 90:10 to 50:50) followed by prep TLC (hexane:ethyl acetate = 50:50) to yield the title compound as a white solid (116 mg, 53%). ^1^H NMR (500 MHz, CDCl_3_) δ 8.02 (d, *J* = 8.5 Hz, 2H), 7.85 – 7.80 (m, 1H), 7.72 – 7.61 (m, 5H), 7.56 (d, *J* = 8.4 Hz, 2H), 7.53 – 7.46 (m, 3H), 7.31 (d, *J* = 8.2 Hz, 2H), 5.55 – 5.31 (m, 1H), 5.22 (s, 2H), 4.85 – 4.78 (m, 2H), 3.62 (pd, *J* = 9.4, 7.3 Hz, 2H), 3.00 – 2.92 (m, 1H), 2.79 – 2.46 (m, 2H), 2.03 – 1.94 (m, 1H), 1.60 (s, 9H), 0.95 (t, *J* = 8.2 Hz, 2H), 0.00 (s, 9H). ^13^C NMR (126 MHz, CDCl_3_) δ 170.79, 168.21, 166.38, 165.71, 165.26, 148.10, 144.35, 140.07, 135.45, 135.07, 134.78, 134.02, 133.28, 132.31, 131.74, 131.13, 131.02, 130.07, 129.66, 128.31, 127.61, 126.91, 126.88, 124.36, 124.18, 81.23, 69.39, 67.64, 55.27, 50.15, 32.08, 28.35, 21.72, 18.27, -1.26. QToF HRMS m/z: calcd for C43H46N4NaO11SSi^+^ [M+Na^+^] = 877.2545; Found 877.2547.

#### *tert*-butyl 4’‘-(((2-(2,6-dioxo-1-((2-(trimethylsilyl)ethoxy)methyl)piperidin-3-yl)-1,3-dioxoisoindolin-4-yl)amino)methyl)-[1,1’‘-biphenyl]-4-carboxylate (11)

Cs_2_CO_3_ (133 mg, 0.407 mmol) and 4-bromothiophenol (51.3 mg, 0.271 mmol) were added to a stirred solution of **10** (116 mg, 0.136 mmol) in DMF (1.36 mL) at room temperature. After 1 hour, the mixture was poured into H_2_O, and extracted with CH_2_Cl_2_. The combined organic layer was dried over anhydrous Na_2_SO_4_ and concentrated. Purification by silica gel column chromatography (hexane:ethyl acetate = 90:10 to 70:30) gave the title compound as a yellow amorphous (81.2 mg, 89%).^1^H NMR (500 MHz, CDCl_3_) δ 8.05 (d, *J* = 8.4 Hz, 2H), 7.61 (dd, *J* = 8.2, 6.0 Hz, 4H), 7.49 – 7.41 (m, 3H), 7.13 (d, *J* = 7.1 Hz, 1H), 6.85 (d, *J* = 8.5 Hz, 1H), 6.74 (t, *J* = 5.9 Hz, 1H), 5.28 (s, 2H), 4.99 – 4.92 (m, 1H), 4.56 (d, *J* = 5.7 Hz, 2H), 3.63 (dtd, *J* = 31.5, 9.8, 6.7 Hz, 2H), 3.03 – 2.94 (m, 1H), 2.86 – 2.73 (m, 2H), 2.17 – 2.07 (m, 1H), 1.61 (s, 9H), 0.95 (ddd, *J* = 9.8, 6.7, 2.7 Hz, 2H), -0.01 (s, 9H). ^13^C NMR (126 MHz, CDCl_3_) δ 171.11, 169.66, 169.23, 167.70, 165.74, 146.70, 144.61, 139.71, 137.74, 136.28, 132.66, 131.06, 130.12, 127.88, 127.64, 126.90, 117.18, 112.21, 110.86, 81.20, 69.30, 67.53, 49.78, 46.61, 32.20, 28.36, 22.15, 18.22, -1.31. QToF HRMS m/z: calcd for C37H43N3NaO7Si^+^ [M+Na^+^] = 692.2762; Found 692.2765.

#### 4’‘-(((2-(2,6-dioxopiperidin-3-yl)-1,3-dioxoisoindolin-4-yl)amino)methyl)-*N*-((1*R*,3*R*)-3-((5-propylpyrazolo[1,5-*a*]pyrimidin-7-yl)amino)cyclopentyl)-[1,1’‘-biphenyl]-4-carboxamide (KI-CDK9d-08)

A mixture of **11** (81.2 mg, 0.121 mmol) and 4 N HCl in 1,4-dioxane (1.21 mL) was stirred at 40 °C overnight. Concentration gave a crude mixture including 4’‘-(((2-(1-(hydroxymethyl)-2,6-dioxopiperidin-3-yl)-1,3-dioxoisoindolin-4-yl)amino)methyl)-[1,1’‘-biphenyl]-4-carboxylic acid (**12**), which was used in the next step without further purification. To a stirred solution of the crude material and KI-ARv3 hydrochloride (61.6 mg, 0.208 mmol) in DMF (1.04 mL), *i*Pr_2_NEt (54.4 mL, 0.313 mmol) and HATU (79.2 mg, 0.208 mmol) were added at room temperature. After stirring overnight, the mixture was poured into H_2_O, and extracted with CH_2_Cl_2_. The combined organic layer was dried over anhydrous Na_2_SO_4_. Filtration and concentration gave a crude mixture, which was purified by silica gel column chromatography (CH_2_Cl_2_:MeOH = 99:1 to 95:5) to yield a mixture of the title compound and 4’‘-(((2-(1- (hydroxymethyl)-2,6-dioxopiperidin-3-yl)-1,3-dioxoisoindolin-4-yl)amino)methyl)-*N*-((1*R*,3*R*)-3-((5-propylpyrazolo[1,5-*a*]pyrimidin-7-yl)amino)cyclopentyl)-[1,1’‘-biphenyl]-4-carboxamide. To a stirred solution of the mixture in DMF (1 mL) was added N1,N2-Dimethylethane-1,2-diamine (9.20 mL, 0.0853 mmol) at 0 °C. After 1.5 hours, the mixture was poured into H_2_O, and extracted with 10% MeOH in CH_2_Cl_2_. The combined organic layer was dried over anhydrous Na_2_SO_4_. Filtration and concentration gave a crude material, which was purified on HPLC (MeCN:H_2_O = 10:90 to 90:10, including 0.1% HCO_2_H) to yield the title compound as a yellow solid (32.2 mg, 52%, 3 steps). ^1^H NMR (500 MHz, DMSO) δ 11.12 (s, 1H), 8.44 (d, *J* = 7.2 Hz, 1H), 8.02 (d, *J* = 2.2 Hz, 1H), 7.94 (d, *J* = 8.5 Hz, 2H), 7.75 (d, *J* = 8.1 Hz, 2H), 7.71 (d, *J* = 7.5 Hz, 2H), 7.55 – 7.46 (m, 3H), 7.30 (t, *J* = 6.3 Hz, 1H), 7.03 (d, *J* = 7.1 Hz, 1H), 6.99 (d, *J* = 8.6 Hz, 1H), 6.30 (d, *J* = 2.1 Hz, 1H), 6.06 (s, 1H), 5.09 (dd, *J* = 12.9, 5.2 Hz, 1H), 4.62 (d, *J* = 6.3 Hz, 2H), 4.51 (q, *J* = 7.2 Hz, 1H), 4.28 (q, *J* = 7.1 Hz, 1H), 2.90 (ddd, *J* = 17.9, 13.8, 5.4 Hz, 1H), 2.65 – 2.52 (m, 5H), 2.30 – 2.21 (m, 1H), 2.20 – 2.01 (m, 3H), 1.85 – 1.59 (m, 4H), 0.92 (t, *J* = 7.4 Hz, 3H). ^13^C NMR (126 MHz, DMSO) δ 172.89, 170.16, 168.81, 167.33, 165.66, 162.20, 148.65, 146.13, 146.06, 143.17, 142.30, 138.98, 137.96, 136.17, 133.47, 132.27, 128.02, 127.68, 127.06, 126.30, 117.71, 110.84, 109.67, 93.64, 85.03, 54.94, 51.54, 49.28, 48.62, 45.13, 38.42, 31.08, 31.03, 30.63, 22.20, 21.96, 13.79. QToF HRMS m/z: calcd for C41H41N8O5^+^ [M+H^+^] = 725.3194; Found 725.3206.

### 4-((4-(2-(2,6-dioxopiperidin-3-yl)-1-oxoisoindolin-5-yl)piperazin-1-yl)methyl)-N-((1R,3R)-3-((5-propylpyrazolo[1,5-a]pyrimidin-7-yl)amino)cyclopentyl)benzamide (KI-CDK9d-32)

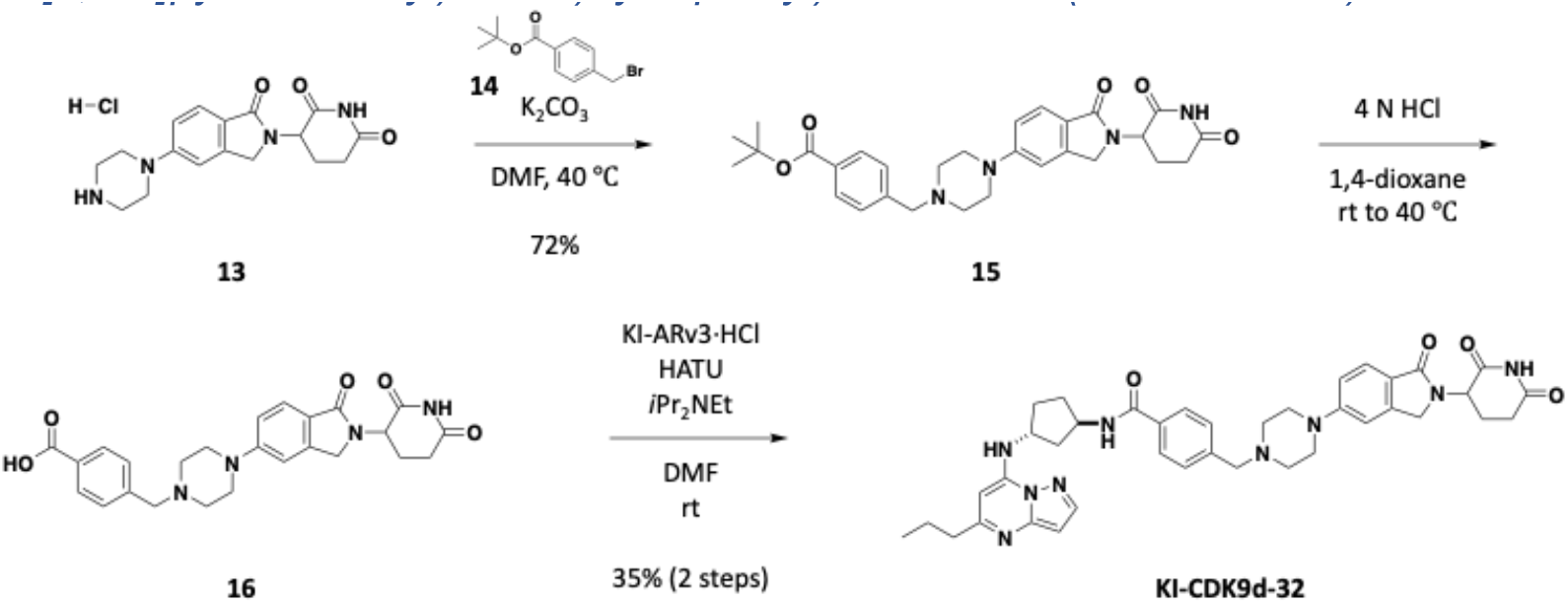

#### *tert*-butyl 4-((4-(2-(2,6-dioxopiperidin-3-yl)-1-oxoisoindolin-5-yl)piperazin-1-yl)methyl)benzoate (15)

*Tert*-butyl 4-(bromomethyl)benzoate (**14**, 7.7 mg, 0.0285 mmol) and K_2_CO_3_ (11.2 mg, 0.0813 mmol) were added to a stirred mixture of 3-(1-oxo-5-(piperazin-1-yl)isoindolin-2-yl)piperidine-2,6-dione^74^ (**13**, 9.9 mg, 0.0271 mmol) in DMF (0.500 mL), and the mixture was warmed to 40 °C. After stirring overnight, the mixture was poured into H_2_O, and extracted with ethyl acetate. The combined organic layer was dried over anhydrous Na_2_SO_4_. Filtration and concentration gave a crude mixture, which was purified by prepTLC (5% methanol in CH_2_Cl_2_) to yield the title compound as a white solid (10.1 mg, 72%). ^1^H NMR (500 MHz, CDCl_3_) δ 8.11 – 8.02 (m, 1H), 8.00 – 7.93 (m, 2H), 7.72 (d, *J* = 8.6 Hz, 1H), 7.41 (d, *J* = 8.2 Hz, 2H), 6.98 (dd, *J* = 8.6, 2.2 Hz, 1H), 6.86 (d, *J* = 2.2 Hz, 1H), 5.19 (dd, *J* = 13.2, 5.0 Hz, 1H), 4.40 (d, *J* = 15.7 Hz, 1H), 4.25 (d, *J* = 15.6 Hz, 1H), 3.61 (s, 2H), 3.32 (t, *J* = 5.0 Hz, 4H), 2.97 – 2.76 (m, 2H), 2.60 (t, *J* = 5.1 Hz, 4H), 2.38 – 2.25 (m, 1H), 2.19 (dtd, *J* = 13.0, 5.3, 2.5 Hz, 1H), 1.60 (s, 9H). ^13^C NMR (126 MHz, CDCl_3_) δ 171.28, 169.88, 169.66, 165.82, 154.56, 143.77, 142.81, 131.24, 129.64, 128.96, 125.15, 121.89, 115.62, 108.39, 81.12, 62.71, 52.93, 51.84, 48.45, 47.03, 31.75, 28.35, 23.63. QToF HRMS m/z: calcd for C29H35N4O5^+^ [M+H^+^] = 519.2602; Found 519.2609.

#### 4-((4-(2-(2,6-dioxopiperidin-3-yl)-1-oxoisoindolin-5-yl)piperazin-1-yl)methyl)-*N*-((1*R*,3*R*)-3-((5-propylpyrazolo[1,5-*a*]pyrimidin-7-yl)amino)cyclopentyl)benzamide (KI-CDK9d-32)

A mixture of **15** (23.9 mg, 0.0461 mmol) and 4 N HCl in 1,4-dioxane (1.00 mL) was stirred at 40 °C overnight. Concentration gave a crude mixture, which was used in the next step without further purification. *i*Pr_2_NEt (24.1 mL, 0.138 mmol) and HATU (35.1 mg, 0.0922 mmol) were added to a stirred solution of the crude material and KI-ARv3 hydrochloride (27.3 mg, 0.0922 mmol) in DMF (1.00 mL) at room temperature. After stirring overnight, the mixture was poured into H_2_O, and extracted with CH_2_Cl_2_. The combined organic layer was dried over anhydrous Na_2_SO_4_. Filtration and concentration gave a crude mixture, which was purified on HPLC (MeCN:H_2_O = 10:90 to 90:10, including 0.1% HCO_2_H) to yield a formic acid salt of the title compound as a white solid (11.2 mg, 35% in 2 steps). ^1^H NMR (500 MHz, DMSO) δ 10.94 (s, 1H), 8.37 (d, *J* = 7.3 Hz, 1H), 8.15 (s, 1H), 8.01 (d, *J* = 2.3 Hz, 1H), 7.83 (d, *J* = 8.0 Hz, 2H), 7.67 (d, *J* = 7.7 Hz, 1H), 7.52 (d, *J* = 8.4 Hz, 1H), 7.43 (d, *J* = 8.0 Hz, 2H), 7.08 – 7.02 (m, 2H), 6.30 (d, *J* = 2.2 Hz, 1H), 6.05 (s, 1H), 5.04 (dd, *J* = 13.3, 5.1 Hz, 1H), 4.54 – 4.43 (m, 1H), 4.32 (d, *J* = 16.9 Hz, 1H), 4.29 – 4.24 (m, 1H), 4.20 (d, *J* = 17.0 Hz, 1H), 3.59 (s, 2H), 3.33 – 3.27 (m, 4H), 2.95 – 2.84 (m, 1H), 2.65 – 2.58 (m, 3H), 2.57 – 2.46 (m, 5H), 2.41 – 2.29 (m, 1H), 2.29 – 2.20 (m, 1H), 2.19 – 1.99 (m, 2H), 1.99 – 1.91 (m, 1H), 1.83 – 1.58 (m, 4H), 0.92 (t, *J* = 7.4 Hz, 3H). ^13^C NMR (126 MHz, DMSO) δ 172.95, 171.31, 168.35, 165.87, 163.22, 162.27, 153.74, 148.83, 146.05, 144.04, 143.10, 141.22, 133.60, 128.64, 127.34, 123.76, 121.53, 114.74, 108.41, 93.68, 84.97, 61.53, 52.34, 51.47, 51.41, 49.19, 47.60, 46.99, 40.43, 38.40, 31.27, 31.06, 30.60, 22.59, 21.95, 13.79. QToF HRMS m/z: calcd for C39H46N9O4^+^ [M+H^+^] = 704.3667; Found 704.3682.

### 4-((4-(2-(1-methyl-2,6-dioxopiperidin-3-yl)-1-oxoisoindolin-5-yl)piperazin-1-yl)methyl)-N-((1R,3R)-3-((5-propylpyrazolo[1,5-a]pyrimidin-7-yl)amino)cyclopentyl)benzamide (KI-CDK9d -D32N)

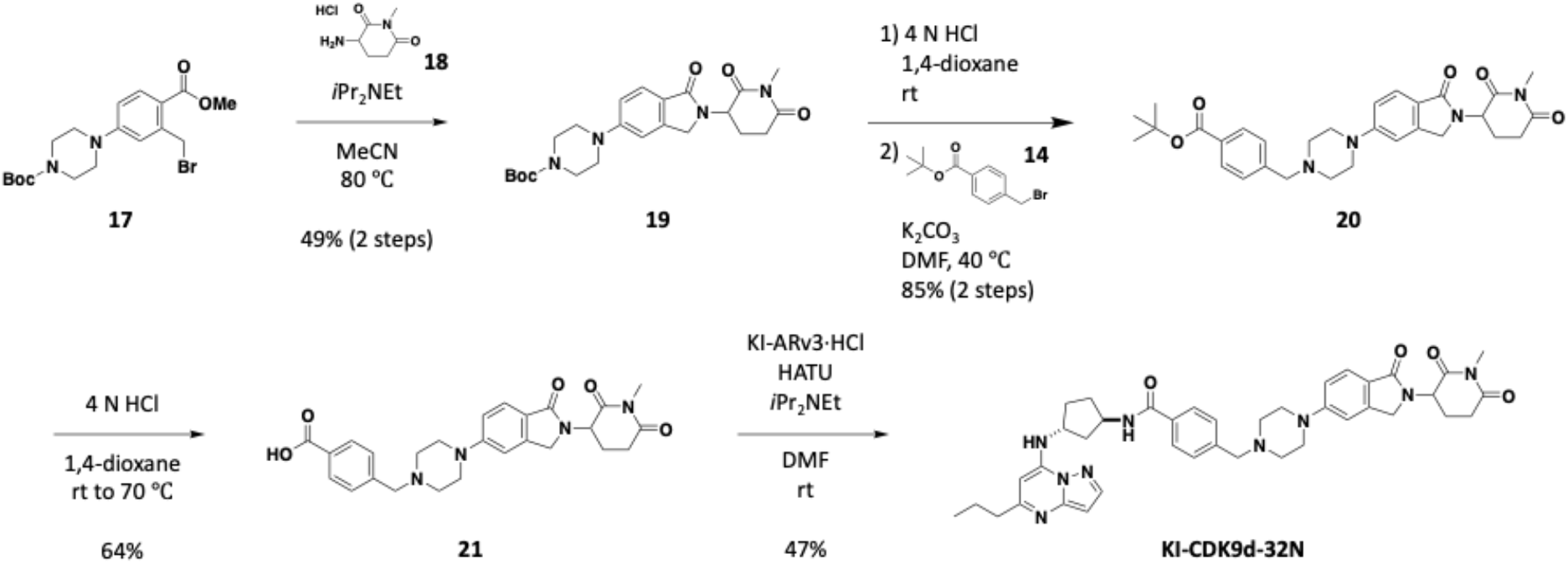

#### *tert*-butyl 4-(2-(1-methyl-2,6-dioxopiperidin-3-yl)-1-oxoisoindolin-5-yl)piperazine-1-carboxylate (19)

A mixture of *tert*-butyl 4-(3-(bromomethyl)-4-(methoxycarbonyl)phenyl)piperazine-1-carboxylate^74^ (**17**, 77.9 mg, 0.188 mmol), 3-amino-1-methylpiperidine-2,6-dione hydrochloride^74^ (**18**, 35.3 mg, 0.198 mmol) and *i*Pr_2_NEt (24.1 mL, 0.138 mmol) in acetonitrile (0.627 mL) was stirred at 80 °C for 24 hours. The mixture was then concentrated and purified by silica gel column chromatography (hexane:ethyl acetate = 50:50 to 0:100) to yield the title compound as a purple solid (40.7 mg, 49%). ^1^H NMR (500 MHz, CDCl_3_) δ 7.75 (d, *J* = 8.7 Hz, 1H), 6.98 (d, *J* = 8.7 Hz, 1H), 6.87 (s, 1H), 5.15 (dd, *J* = 13.5, 5.0 Hz, 1H), 4.38 (d, *J* = 15.4 Hz, 1H), 4.25 (d, *J* = 15.6 Hz, 1H), 3.59 (t, *J* = 5.2 Hz, 4H), 3.27 (t, *J* = 5.2 Hz, 4H), 3.17 (s, 3H), 3.02 – 2.93 (m, 1H), 2.90 – 2.79 (m, 1H), 2.35 – 2.22 (m, 1H), 2.20 – 2.11 (m, 1H), 1.48 (s, 9H). ^13^C NMR (126 MHz, CDCl_3_) δ 171.50, 170.48, 169.61, 154.78, 154.42, 143.82, 125.22, 122.72, 116.06, 108.91, 80.29, 76.91, 52.52, 48.62, 47.20, 32.24, 28.55, 27.28, 22.96. QToF HRMS m/z: calcd for C23H31N4O5^+^ [M+H^+^] = 443.2289; Found 443.2296.

#### *tert*-butyl 4-((4-(2-(1-methyl-2,6-dioxopiperidin-3-yl)-1-oxoisoindolin-5-yl)piperazin-1-yl)methyl)benzoate (20)

A mixture of **19** (40.7 mg, 0.0920 mmol) and 4 N HCl in 1,4-dioxane (1.00 mL) was stirred at room temperature. After 3 hours, the mixture was concentrated to give a crude mixture, which was used in the next reaction without further purification. *Tert*-butyl 4-(bromomethyl)benzoate (**14**, 26.2 mg, 0.0966 mmol) and K_2_CO_3_ (38.1 mg, 0.276 mmol) were added to a stirred solution of the crude mixture in DMF (1.00 mL), then the mixture was warmed to 40 °C. After stirring overnight, the mixture was poured into H_2_O, and extracted with ethyl acetate. The combined organic layer was dried over anhydrous Na_2_SO_4_. Filtration and concentration gave a crude mixture, which was purified by silica gel column chromatography (hexane:ethyl acetate = 50:50 to 0:100) to yield the title compound as a white foam (41.8 mg, 85% in 2 steps). ^1^H NMR (500 MHz, CDCl_3_) δ 7.96 (d, *J* = 7.8 Hz, 2H), 7.73 (d, *J* = 8.6 Hz, 1H), 7.41 (d, *J* = 8.0 Hz, 2H), 6.98 (d, *J* = 8.5 Hz, 1H), 6.86 (s, 1H), 5.15 (dd, *J* = 13.4, 5.0 Hz, 1H), 4.37 (d, *J* = 15.6 Hz, 1H), 4.24 (d, *J* = 15.6 Hz, 1H), 3.61 (s, 2H), 3.31 (t, *J* = 5.0 Hz, 4H), 3.17 (s, 3H), 3.02 – 2.93 (m, 1H), 2.90 – 2.79 (m, 1H), 2.60 (t, *J* = 5.0 Hz, 4H), 2.27 (tt, *J* = 13.3, 6.7 Hz, 1H), 2.19 – 2.10 (m, 1H), 1.59 (s, 9H). ^13^C NMR (126 MHz, CDCl_3_) δ 171.54, 170.52, 169.73, 165.81, 154.52, 143.80, 142.85, 131.21, 129.63, 128.94, 125.11, 122.15, 115.62, 108.41, 81.10, 62.72, 52.95, 52.50, 48.50, 47.20, 32.26, 28.35, 27.27, 22.97. QToF HRMS m/z: calcd for C30H37N4O5^+^ [M+H^+^] = 533.2758; Found 533.2762.

#### 4-((4-(2-(1-methyl-2,6-dioxopiperidin-3-yl)-1-oxoisoindolin-5-yl)piperazin-1-yl)methyl)benzoic acid (21)

A mixture of **20** (41.8 mg, 0.0785 mmol) and 4 N HCl in 1,4-dioxane (2.00 mL) was warmed to 40 °C. After stirring overnight, the mixture was heated to 70 °C and stirred for 2 hours. The mixture was then concentrated and purified on HPLC (MeCN:H_2_O = 10:90 to 90:10, including 0.1% HCO_2_H) to yield the title compound as a white solid (23.8 mg, 64%). ^1^H NMR (500 MHz, Pyr) δ 8.49 (d, *J* = 7.9 Hz, 2H), 7.98 (d, *J* = 8.5 Hz, 1H), 7.66 – 7.55 (m, 2H), 7.08 (dd, *J* = 8.5, 2.2 Hz, 1H), 7.00 (d, *J* = 2.2 Hz, 1H), 5.59 (dd, *J* = 13.7, 5.0 Hz, 1H), 4.51 (d, *J* = 16.1 Hz, 1H), 4.39 (d, *J* = 16.0 Hz, 1H), 3.58 (s, 2H), 3.36 – 3.30 (m, 4H), 3.12 (s, 3H), 2.96 – 2.89 (m, 2H), 2.57 (t, *J* = 5.0 Hz, 4H), 2.44 – 2.33 (m, 1H), 2.08 – 1.99 (m, 1H). ^13^C NMR (126 MHz, Pyr) δ 172.19, 171.63, 170.02, 169.41, 155.01, 145.01, 144.11, 131.90, 130.71, 129.65, 125.15, 123.16, 115.91, 109.31, 62.90, 53.44, 53.29, 48.80, 47.88, 32.71, 27.26, 23.27. QToF HRMS m/z: calcd for C26H29N4O5^+^ [M+H^+^] = 477.2132; Found 477.2142.

#### 4-((4-(2-(1-methyl-2,6-dioxopiperidin-3-yl)-1-oxoisoindolin-5-yl)piperazin-1-yl)methyl)-*N*-((1*R*,3*R*)-3-((5-propylpyrazolo[1,5-*a*]pyrimidin-7-yl)amino)cyclopentyl)benzamide (KI-CDK9d-32N)

*i*Pr_2_NEt (26.1 mL, 0.150 mmol) and HATU (57.0 mg, 0.150 mmol) were added to a stirred solution of **21** (23.8 mg, 0.0499 mmol) and KI-ARv3 hydrochloride (29.5 mg, 0.0999 mmol) in DMF (1.00 mL) at room temperature. After stirring overnight, the mixture was poured into H_2_O, and extracted with 10% methanol in CH_2_Cl_2_. The combined organic layer was dried over anhydrous Na_2_SO_4_. Filtration and concentration gave a crude mixture, which was purified on HPLC (MeCN:H_2_O = 10:90 to 90:10, including 0.1% HCO_2_H) and prepTLC (8% methanol in CH_2_Cl_2_) to yield the title compound as a white solid (16.9 mg, 47%). ^1^H NMR (500 MHz, DMSO) δ 8.32 (d, *J* = 7.3 Hz, 1H), 7.97 (d, *J* = 2.3 Hz, 1H), 7.79 (d, *J* = 7.9 Hz, 2H), 7.63 (d, *J* = 7.6 Hz, 1H), 7.49 (d, *J* = 8.8 Hz, 1H), 7.39 (d, *J* = 8.0 Hz, 2H), 7.04 – 6.98 (m, 2H), 6.26 (d, *J* = 2.2 Hz, 1H), 6.00 (s, 1H), 5.07 (dd, *J* = 13.5, 5.0 Hz, 1H), 4.45 (q, *J* = 7.1 Hz, 1H), 4.28 (d, *J* = 16.9 Hz, 1H), 4.23 (q, *J* = 7.1 Hz, 1H), 4.15 (d, *J* = 17.0 Hz, 1H), 3.54 (s, 2H), 3.26 (t, *J* = 4.8 Hz, 3H), 2.95 (s, 3H), 2.98 – 2.88 (m, 1H), 2.75 – 2.66 (m, 1H), 2.61 – 2.54 (m, 2H), 2.49 – 2.44 (m, 4H), 2.37 – 2.26 (m, 1H), 2.25 – 2.16 (m, 1H), 2.15 – 1.98 (m, 3H), 1.98 – 1.88 (m, 1H), 1.79 – 1.55 (m, 5H), 0.88 (t, *J* = 7.4 Hz, 3H). ^13^C NMR (126 MHz, DMSO) δ 171.97, 170.94, 168.37, 165.86, 162.26, 153.76, 148.83, 146.05, 144.05, 143.09, 141.24, 133.59, 128.62, 127.33, 123.79, 121.51, 114.73, 108.39, 93.67, 84.96, 61.53, 52.34, 51.95, 51.47, 49.19, 47.60, 47.01, 40.43, 38.39, 31.42, 31.05, 30.60, 26.56, 21.94, 21.83, 13.79. QToF HRMS m/z: calcd for C40H48N9O4^+^ [M+H^+^] = 718.3824; Found 718.3834.

## References

1. Agudo-Ibáñez, L., Morante, M., García-Gutiérrez, L., Quintanilla, A., Rodríguez, J., Muñoz, A., León, J., and Crespo, P. (2023). ERK2 stimulates MYC transcription by anchoring CDK9 to the MYC promoter in a kinase activity–independent manner. Sci. Signal. 16. 10.1126/SCISIGNAL.ADG4193.

2. Lu, H., Xue, Y., Yu, G.K., Arias, C., Lin, J., Fong, S., Faure, M., Weisburd, B., Ji, X., Mercier, A., et al. (2015). Compensatory induction of MYC expression by sustained CDK9 inhibition via a BRD4-dependent mechanism. Elife 4. 10.7554/ELIFE.06535.

3. Meyer, N., and Penn, L.Z. (2008). Reflecting on 25 years with MYC. Nat. Rev. Cancer 2008 812 8, 976–990. 10.1038/nrc2231.

4. Blake, D.R., Vaseva, A. V., Hodge, R.G., Kline, M.P., Gilbert, T.S.K., Tyagi, V., Huang, D., Whiten, G.C., Larson, J.E., Wang, X., et al. (2019). Application of a MYC degradation screen identifies sensitivity to CDK9 inhibitors in KRAS-mutant pancreatic cancer. Sci. Signal. 12, 7259. 10.1126/SCISIGNAL.AAV7259.

5. Gargano, B., Amente, S., Majello, B., and Lania, L. (2007). P-TEFb is a crucial co-factor for Myc transactivation. Cell Cycle 6, 2031–2037. 10.4161/cc.6.16.4554.

6. Toure, M.A., and Koehler, A.N. (2023). Addressing Transcriptional Dysregulation in Cancer through CDK9 Inhibition. Biochemistry 62, 1114–1123. 10.1021/ACS.BIOCHEM.2C00609/ASSET/IMAGES/LARGE/BI2C00609_0002.JPEG.

7. Chirnomas, D., Hornberger, K.R., and Crews, C.M. (2023). Protein degraders enter the clinic — a new approach to cancer therapy. Nat. Rev. Clin. Oncol. 2023 204 20, 265–278. 10.1038/s41571-023-00736-3.

8. Pettersson, M., and Crews, C.M. (2019). PROteolysis TArgeting Chimeras (PROTACs) — Past, present and future. Drug Discov. Today Technol. 31, 15–27. 10.1016/j.ddtec.2019.01.002.

9. Gao, X., III, H.A.B. Vuky, J., Dreicer, R., Sartor, A.O., Sternberg, C.N., Percent, I.J., Hussain, M.H.A., Kalebasty, A.R., Shen, J., et al. (2022). Phase 1/2 study of ARV-110, an androgen receptor (AR) PROTAC degrader, in metastatic castration-resistant prostate cancer (mCRPC). 10.1200/JCO.2022.40.6_suppl.017 40, 17–17. 10.1200/JCO.2022.40.6_SUPPL.017.

10. Olson, C.M., Jiang, B., Erb, M.A., Liang, Y., Doctor, Z.M., Zhang, Z., Zhang, T., Kwiatkowski, N., Boukhali, M., Green, J.L., et al. (2018). Pharmacological perturbation of CDK9 using selective CDK9 inhibition or degradation. Nat. Chem. Biol. 14, 163–170. 10.1038/nchembio.2538.

11. Li, J., Liu, T., Song, Y., Wang, M., Liu, L., Zhu, H., Li, Q., Lin, J., Jiang, H., Chen, K., et al. (2022). Discovery of Small-Molecule Degraders of the CDK9-Cyclin T1 Complex for Targeting Transcriptional Addiction in Prostate Cancer. J. Med. Chem. 65, 11034–11057. 10.1021/ACS.JMEDCHEM.2C00257/SUPPL_FILE/JM2C00257_SI_006.PDB.

12. King, H.M., Rana, S., Kubica, S.P., Mallareddy, J.R., Kizhake, S., Ezell, E.L., Zahid, M., Naldrett, M.J., Alvarez, S., Law, H.C.H., et al. (2021). Aminopyrazole based CDK9 PROTAC sensitizes pancreatic cancer cells to venetoclax. Bioorganic Med. Chem. Lett. 43. 10.1016/J.BMCL.2021.128061.

13. Richters, A., Doyle, S.K., Freeman, D.B., Lee, C., Leifer, B.S., Jagannathan, S., Kabinger, F., Koren, J.V., Struntz, N.B., Urgiles, J., et al. (2020). Modulating Androgen Receptor-Driven Transcription in Prostate Cancer with Selective CDK9 Inhibitors. Cell Chem. Biol. 28. 10.1016/j.chembiol.2020.10.001.

14. Freeman, D.B., Hopkins, T.D., Mikochik, P.J., Vacca, J.P., Gao, H., Naylor-Olsen, A., Rudra, S., Li, H., Pop, M.S., Villagomez, R.A., et al. (2023). Discovery of KB-0742, a Potent, Selective, Orally Bioavailable Small Molecule Inhibitor of CDK9 for MYC-Dependent Cancers. J. Med. Chem. 66, 15629–15647. 10.1021/ACS.JMEDCHEM.3C01233/SUPPL_FILE/JM3C01233_SI_001.PDF.

15. Day, M.A., Saffran, D.C., Hood, T., Obholzer, N., Pandey, A., Lin, C.Y., Kumar, P., Freed, D.M., and DiMartino, J. (2022). Abstract 2564: CDK9 inhibition via KB-0742 is a potential strategy to treat transcriptionally addicted cancers. Cancer Res. 82, 2564–2564. 10.1158/1538-7445.AM2022-2564.

16. Saffran, D.C., Day, M.A.L., Rioux, N., Chen, T., Lee, C., Amara, S.N.-A., Freeman, D.B., Hood, T., Lin, C.Y., Kumar, P., et al. (2022). Abstract P5-08-05: Preclinical activity of KB-0742, an oral, highly selective, CDK9 inhibitor, in cell lines and in MYC-high expressing, patient-derived models of multiple breast cancer subtypes. Cancer Res. 82, P5-08–05. 10.1158/1538-7445.SABCS21-P5-08-05.

17. Day, M.A.L., Saffran, D.C., Rioux, N., Chen, T., Lee, C., Freeman, D.B., MacKenzie, C., Vacca, J.P., Rahl, P.B., Trotter, B.W., et al. (2021). Abstract P228: Preclinical pharmacokinetics and pharmacodynamics of KB-0742, a selective, oral CDK9 inhibitor. Mol. Cancer Ther. 20, P228–P228. 10.1158/1535-7163.TARG-21-P228.

18. Bradner, J.E., McPherson, O.M., Mazitschek, R., Barnes-Seeman, D., Shen, J.P., Dhaliwal, J., Stevenson, K.E., Duffner, J.L., Park, S.B., Neuberg, D.S., et al. (2006). A Robust Small-Molecule Microarray Platform for Screening Cell Lysates. Chem. Biol. 13, 493–504. 10.1016/J.CHEMBIOL.2006.03.004.

19. Ito, T., Ando, H., Suzuki, T., Ogura, T., Hotta, K., Imamura, Y., Yamaguchi, Y., and Handa, H. (2010). Identification of a primary target of thalidomide teratogenicity. Science 327, 1345–1350. 10.1126/SCIENCE.1177319.

20. Békés, M., Langley, D.R., and Crews, C.M. (2022). PROTAC targeted protein degraders: the past is prologue. Nat. Rev. Drug Discov. 2022 213 21, 181–200. 10.1038/s41573-021-00371-6.

21. Riching, K.M., Mahan, S., Corona, C.R., McDougall, M., Vasta, J.D., Robers, M.B., Urh, M., and Daniels, D.L. (2018). Quantitative Live-Cell Kinetic Degradation and Mechanistic Profiling of PROTAC Mode of Action. ACS Chem. Biol. 13, 2758–2770. 10.1021/ACSCHEMBIO.8B00692.

22. Riching, K.M., Caine, E.A., Urh, M., and Daniels, D.L. (2022). The importance of cellular degradation kinetics for understanding mechanisms in targeted protein degradation. Chem. Soc. Rev. 51, 6210–6221. 10.1039/D2CS00339B.

23. Zografou-Barredo, N.A., Hallatt, A.J., Goujon-Ricci, J., and Cano, C. (2023). A beginner’‘s guide to current synthetic linker strategies towards VHL-recruiting PROTACs. Bioorg. Med. Chem. 88–89, 117334. 10.1016/J.BMC.2023.117334.

24. Nguyen, T.M., Sreekanth, V., Deb, A., Kokkonda, P., Tiwari, P.K., Donovan, K.A., Shoba, V., Chaudhary, S.K., Mercer, J.A.M., Lai, S., et al. (2023). Proteolysis Targeting Chimeras With Reduced Off-targets. bioRxiv, 2021.11.18.468552. 10.1101/2021.11.18.468552.

25. Haid, R.T.U., and Reichel, A. (2023). A Mechanistic Pharmacodynamic Modeling Framework for the Assessment and Optimization of Proteolysis Targeting Chimeras (PROTACs). Pharmaceutics 15. 10.3390/PHARMACEUTICS15010195/S1.

26. Schwinn, M.K., Machleidt, T., Zimmerman, K., Eggers, C.T., Dixon, A.S., Hurst, R., Hall, M.P., Encell, L.P., Binkowski, B.F., and Wood, K. V. (2018). CRISPR-Mediated Tagging of Endogenous Proteins with a Luminescent Peptide. ACS Chem. Biol. 13, 467–474. 10.1021/ACSCHEMBIO.7B00549/ASSET/IMAGES/LARGE/CB-2017-005496_0008.JPEG.

27. Thng, D.K.H., Toh, T.B., and Chow, E.K.H. (2021). Capitalizing on Synthetic Lethality of MYC to Treat Cancer in the Digital Age. Trends Pharmacol. Sci. 42, 166–182. 10.1016/J.TIPS.2020.11.014.

28. Chen, H., Liu, H., and Qing, G. (2018). Targeting oncogenic Myc as a strategy for cancer treatment. Signal Transduct. Target. Ther. 2018 31 3, 1–7. 10.1038/s41392-018-0008-7.

29. Liberzon, A., Birger, C., Thorvaldsdóttir, H., Ghandi, M., Mesirov, J.P., and Tamayo, P. (2015). The Molecular Signatures Database (MSigDB) hallmark gene set collection. Cell Syst. 1, 417–425. 10.1016/J.CELS.2015.12.004.

30. Kanehisa, M., and Goto, S. (2000). KEGG: kyoto encyclopedia of genes and genomes. Nucleic Acids Res. 28, 27–30. 10.1093/NAR/28.1.27.

31. Boulon, S., Westman, B.J., Hutten, S., Boisvert, F.M., and Lamond, A.I. (2010). The Nucleolus under Stress. Mol. Cell 40, 216. 10.1016/J.MOLCEL.2010.09.024.

32. Lovly, C.M., and Shaw, A.T. (2014). Molecular Pathways: Resistance to Kinase Inhibitors and Implications for Therapeutic Strategies. Clin. Cancer Res. 20, 2249. 10.1158/1078-0432.CCR-13-1610.

33. Lovén, J., Orlando, D.A., Sigova, A.A., Lin, C.Y., Rahl, P.B., Burge, C.B., Levens, D.L., Lee, T.I., and Young, R.A. (2012). Revisiting global gene expression analysis. Cell 151, 476–482. 10.1016/J.CELL.2012.10.012.

34. Mann, M.W., Fu, Y., Gearhart, R.L., Xu, X., Roberts, D.S., Li, Y., Zhou, J., Ge, Y., and Brasier, A.R. (2023). Bromodomain-containing Protein 4 regulates innate inflammation via modulation of alternative splicing. Front. Immunol. 14, 1212770. 10.3389/FIMMU.2023.1212770/FULL.

35. Egloff, S. (2021). CDK9 keeps RNA polymerase II on track. Cell. Mol. Life Sci. 2021 7814 78, 5543–5567. 10.1007/S00018-021-03878-8.

36. Zheng, B., Gold, S., Iwanaszko, M., Howard, B.C., Wang, L., and Shilatifard, A. (2023). Distinct layers of BRD4-PTEFb reveal bromodomain-independent function in transcriptional regulation. Mol. Cell 83, 2896-2910.e4. 10.1016/j.molcel.2023.06.032.

37. Feng, C., Song, C., Jiang, Y., Zhao, J., Zhang, J., Wang, Y., Yin, M., Zhu, J., Ai, B., Wang, Q., et al. (2023). Landscape and significance of human super enhancer-driven core transcription regulatory circuitry. Mol. Ther. - Nucleic Acids 32, 385–401. 10.1016/j.omtn.2023.03.014.

38. Chen, Y., Xu, L., Lin, R.Y.T., Müschen, M., and Koeffler, H.P. (2020). Core transcriptional regulatory circuitries in cancer. Oncogene 2020 3943 39, 6633–6646. 10.1038/s41388-020-01459-w.

39. Felsher, D.W. (2010). MYC Inactivation Elicits Oncogene Addiction through Both Tumor Cell– Intrinsic and Host-Dependent Mechanisms. Genes Cancer 1, 597. 10.1177/1947601910377798.

40. Van Riggelen, J., Yetil, A., and Felsher, D.W. (2010). MYC as a regulator of ribosome biogenesis and protein synthesis. Nat. Rev. Cancer 2010 104 10, 301–309. 10.1038/nrc2819.

41. Burger, K., Mühl, B., Rohrmoser, M., Coordes, B., Heidemann, M., Kellner, M., Gruber-Eber, A., Heissmeyer, V., Strässer, K., and Eick, D. (2013). Cyclin-dependent Kinase 9 Links RNA Polymerase II Transcription to Processing of Ribosomal RNA. J. Biol. Chem. 288, 21173. 10.1074/JBC.M113.483719.

42. Abraham, K.J., Khosraviani, N., Chan, J.N.Y., Gorthi, A., Samman, A., Zhao, D.Y., Wang, M., Bokros, M., Vidya, E., Ostrowski, L.A., et al. (2020). Nucleolar RNA polymerase II drives ribosome biogenesis. Nat. 2020 5857824 585, 298–302. 10.1038/s41586-020-2497-0.

43. Mitrea, D.M., Cika, J.A., Stanley, C.B., Nourse, A., Onuchic, P.L., Banerjee, P.R., Phillips, A.H., Park, C.G., Deniz, A.A., and Kriwacki, R.W. (2018). Self-interaction of NPM1 modulates multiple mechanisms of liquid–liquid phase separation. Nat. Commun. 9. 10.1038/S41467-018-03255-3.

44. Stenström, L., Mahdessian, D., Gnann, C., Cesnik, A.J., Ouyang, W., Leonetti, M.D., Uhlén, M., Cuylen-Haering, S., Thul, P.J., and Lundberg, E. (2020). Mapping the nucleolar proteome reveals a spatiotemporal organization related to intrinsic protein disorder. Mol. Syst. Biol. 16, 9469. 10.15252/MSB.20209469.

45. Calo, E., Flynn, R.A., Martin, L., Spitale, R.C., Chang, H.Y., and Wysocka, J. (2015). RNA helicase DDX21 coordinates transcription and ribosomal RNA processing. Nature 518, 249. 10.1038/NATURE13923.

46. Liu, H., and Herrmann, C.H. (2005). Differential localization and expression of the Cdk9 42k and 55k isoforms. J. Cell. Physiol. 203, 251–260. 10.1002/JCP.20224.

47. Sridharan, S., Hernandez-Armendariz, A., Kurzawa, N., Potel, C.M., Memon, D., Beltrao, P., Bantscheff, M., Huber, W., Cuylen-Haering, S., and Savitski, M.M. (2022). Systematic discovery of biomolecular condensate-specific protein phosphorylation. Nat. Chem. Biol. 2022 1810 18, 1104–1114. 10.1038/s41589-022-01062-y.

48. Batool, A., Majeed, S.T., Aashaq, S., Majeed, R., and Andrabi, K.I. (2020). Eukaryotic Initiation Factor 4E phosphorylation acts a switch for its binding to 4E-BP1 and mRNA cap assembly. Biochem. Biophys. Res. Commun. 527, 489–495. 10.1016/J.BBRC.2020.04.086.

49. Tang, Y., Luo, J., Yang Liu, S., Zheng, H., Zhan, Y., Fan, S., and Wen, Q. (2022). Overexpression of p-4EBP1 associates with p-eIF4E and predicts poor prognosis for non-small cell lung cancer patients with resection. PLoS One 17. 10.1371/JOURNAL.PONE.0265465.

50. Corsello, S.M., Nagari, R.T., Spangler, R.D., Rossen, J., Kocak, M., Bryan, J.G., Humeidi, R., Peck, D., Wu, X., Tang, A.A., et al. (2020). Discovering the anti-cancer potential of non-oncology drugs bysystematic viability profiling. Nat. cancer 1, 235. 10.1038/S43018-019-0018-6.

51. Chang, L., Ruiz, P., Ito, T., and Sellers, W.R. (2021). Targeting pan-essential genes in cancer: Challenges and opportunities. Cancer Cell 39, 466–479. 10.1016/J.CCELL.2020.12.008.

52. Kurimchak, A.M., Herrera-Montávez, C., Montserrat-Sangrà, S., Araiza-Olivera, D., Hu, J., Neumann-Domer, R., Kuruvilla, M., Bellacosa, A., Testa, J.R., Jin, J., et al. (2022). The drug efflux pump MDR1 promotes intrinsic and acquired resistance to PROTACs in cancer cells HHS Public Access. Sci Signal 15, 2707. 10.1126/scisignal.abn2707.

53. Garnett, M.J., Edelman, E.J., Heidorn, S.J., Greenman, C.D., Dastur, A., Lau, K.W., Greninger, P., Thompson, I.R., Luo, X., Soares, J., et al. (2012). Systematic identification of genomic markers of drug sensitivity in cancer cells. Nature 483. 10.1038/nature11005.

54. Dang, C. V., O’Donnell, K.A., Zeller, K.I., Nguyen, T., Osthus, R.C., and Li, F. (2006). The c-Myc target gene network. Semin. Cancer Biol. 16, 253–264. 10.1016/J.SEMCANCER.2006.07.014.

55. Hann, S.R. (2014). MYC Cofactors: Molecular Switches Controlling Diverse Biological Outcomes. Cold Spring Harb. Perspect. Med. 4. 10.1101/CSHPERSPECT.A014399.

56. Anshabo, A.T., Milne, R., Wang, S., and Albrecht, H. (2021). CDK9: A Comprehensive Review of Its Biology, and Its Role as a Potential Target for Anti-Cancer Agents. Front. Oncol. 11, 1573. 10.3389/fonc.2021.678559.

57. Bacon, C.W., and D’Orso, I. (2019). CDK9: a signaling hub for transcriptional control. Transcription 10, 57–75. 10.1080/21541264.2018.1523668.

58. Dey, J., Deckwerth, T.L., Kerwin, W.S., Casalini, J.R., Merrell, A.J., Grenley, M.O., Burns, C., Ditzler, S.H., Dixon, C.P., Beirne, E., et al. (2017). Voruciclib, a clinical stage oral CDK9 inhibitor, represses MCL-1 and sensitizes high-risk Diffuse Large B-cell Lymphoma to BCL2 inhibition. Sci. Reports 2017 71 7, 1–11. 10.1038/s41598-017-18368-w.

59. Burslem, G.M., Smith, B.E., Lai, A.C., Jaime-Figueroa, S., McQuaid, D.C., Bondeson, D.P., Toure, M., Dong, H., Qian, Y., Wang, J., et al. (2018). The Advantages of Targeted Protein Degradation Over Inhibition: An RTK Case Study. Cell Chem. Biol. 25, 67-77.e3. 10.1016/J.CHEMBIOL.2017.09.009.

60. Wilmarth, P.A., Riviere, M.A., and David, L.L. (2009). Techniques for accurate protein identification in shotgun proteomic studies of human, mouse, bovine, and chicken lenses. J. Ocul. Biol. Dis. Infor. 2, 223–234. 10.1007/S12177-009-9042-6/FIGURES/5.

61. Wilmarth, P. PAW Pipeline: A Comet-based, best practices proteomics pipeline. https://github.com/pwilmart/PAW_pipeline.

62. Taus, T., Köcher, T., Pichler, P., Paschke, C., Schmidt, A., Henrich, C., and Mechtler, K. (2011). Universal and confident phosphorylation site localization using phosphoRS. J. Proteome Res. 10, 5354–5362. 10.1021/PR200611N.

63. Hornbeck, P. V., Zhang, B., Murray, B., Kornhauser, J.M., Latham, V., and Skrzypek, E. (2015). PhosphoSitePlus, 2014: mutations, PTMs and recalibrations. Nucleic Acids Res. 43, D512–D520. 10.1093/NAR/GKU1267.

64. Yu, G., Wang, L.G., Han, Y., and He, Q.Y. (2012). clusterProfiler: an R Package for Comparing Biological Themes Among Gene Clusters. OMICS 16, 284. 10.1089/OMI.2011.0118.

65. Mildrum, S., Hendricks, A., Stortchevoi, A., Kamelamela, N., Butty, V.L., and Levine, S.S. (2020). High-throughput Minitaturized RNA-Seq Library Preparation. J. Biomol. Tech. 31, 151–156. 10.7171/JBT.20-3104-004.

66. Peltzer, A., Mohr, C., Stadermann, K.B., Zwick, M., and Schmid, R. (2024). nf-core/nanostring: a pipeline for reproducible NanoString nCounter analysis. Bioinformatics 40. 10.1093/BIOINFORMATICS/BTAE019.

67. Soneson, C., Love, M.I., and Robinson, M.D. (2016). Differential analyses for RNA-seq: Transcript-level estimates improve gene-level inferences. F1000Research 4. 10.12688/F1000RESEARCH.7563.2/DOI.

68. Wickham, H., Averick, M., Bryan, J., Chang, W. D’ L., Mcgowan, A., François, R., Grolemund, G., Hayes, A., Henry, L., et al. (2019). Welcome to the Tidyverse. J. Open Source Softw. 4, 1686. 10.21105/JOSS.01686.

69. Love, M.I., Huber, W., and Anders, S. (2014). Moderated estimation of fold change and dispersion for RNA-seq data with DESeq2. Genome Biol. 15, 1–21. 10.1186/S13059-014-0550-8/FIGURES/9.

70. Anders, S., and Huber, W. (2010). Differential expression analysis for sequence count data. Genome Biol 11, 106. 10.1186/gb-2010-11-10-r106.

71. Zhu, A., Ibrahim, J.G., and Love, M.I. (2019). Heavy-tailed prior distributions for sequence count data: removing the noise and preserving large differences. Bioinformatics 35, 2084–2092. 10.1093/BIOINFORMATICS/BTY895.

72. Mootha, V.K., Lindgren, C.M., Eriksson, K.F., Subramanian, A., Sihag, S., Lehar, J., Puigserver, P., Carlsson, E., Ridderstråle, M., Laurila, E., et al. (2003). PGC-1alpha-responsive genes involved in oxidative phosphorylation are coordinately downregulated in human diabetes. Nat. Genet. 34, 267–273. 10.1038/NG1180.

73. Subramanian, A., Tamayo, P., Mootha, V.K., Mukherjee, S., Ebert, B.L., Gillette, M.A., Paulovich, A., Pomeroy, S.L., Golub, T.R., Lander, E.S., et al. (2005). From the Cover: Gene set enrichment analysis: A knowledge-based approach for interpreting genome-wide expression profiles. Proc. Natl. Acad. Sci. U. S. A. 102, 15545. 10.1073/PNAS.0506580102.

74. Crews, A.P., Berlin, M., Dong, H., Ishchenko, A., Cacace, A.M., and Chandler, J.T. (2020). Tauprotein targeting compounds and associated methods of use.

