## Supplementary Figures and Tables for "Targeted Degradation of CDK9 Potently Disrupts the MYC Transcriptional Network"

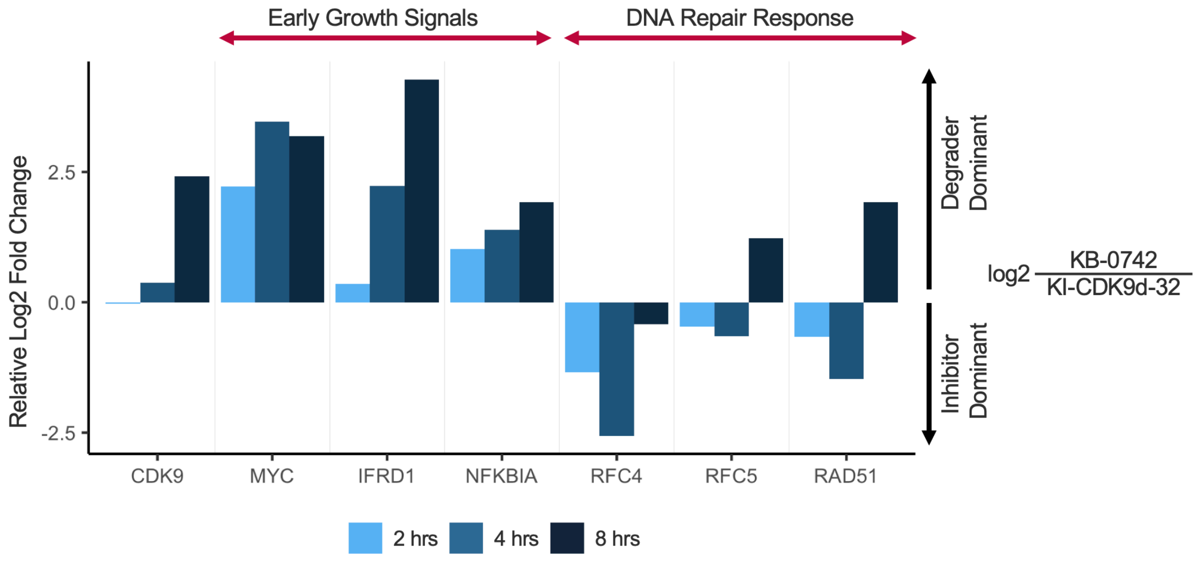


**Figure S3.1: Differential Effects from CDK9 degradation and inhibition**

RNA sequencing evaluation of transcript levels following 2, 4, and 8 hours of treatment with 1.2 μM of KB-0742 and 15nM of KI-CDK9d-32. Barplot showing log2 fold change in mRNA levels of select genes in key cancer pathways that are differentially impacted in a degrader versus inhibitor direct comparison. Genes shown have high statistical significance. The transcripts of genes with LFC > 0 are differentially downregulated by the degrader. See S3.2 and S3.3 for GSEA enrichment plots of MYC target genes, TNFA via NFKB signaling, and DNA repair.

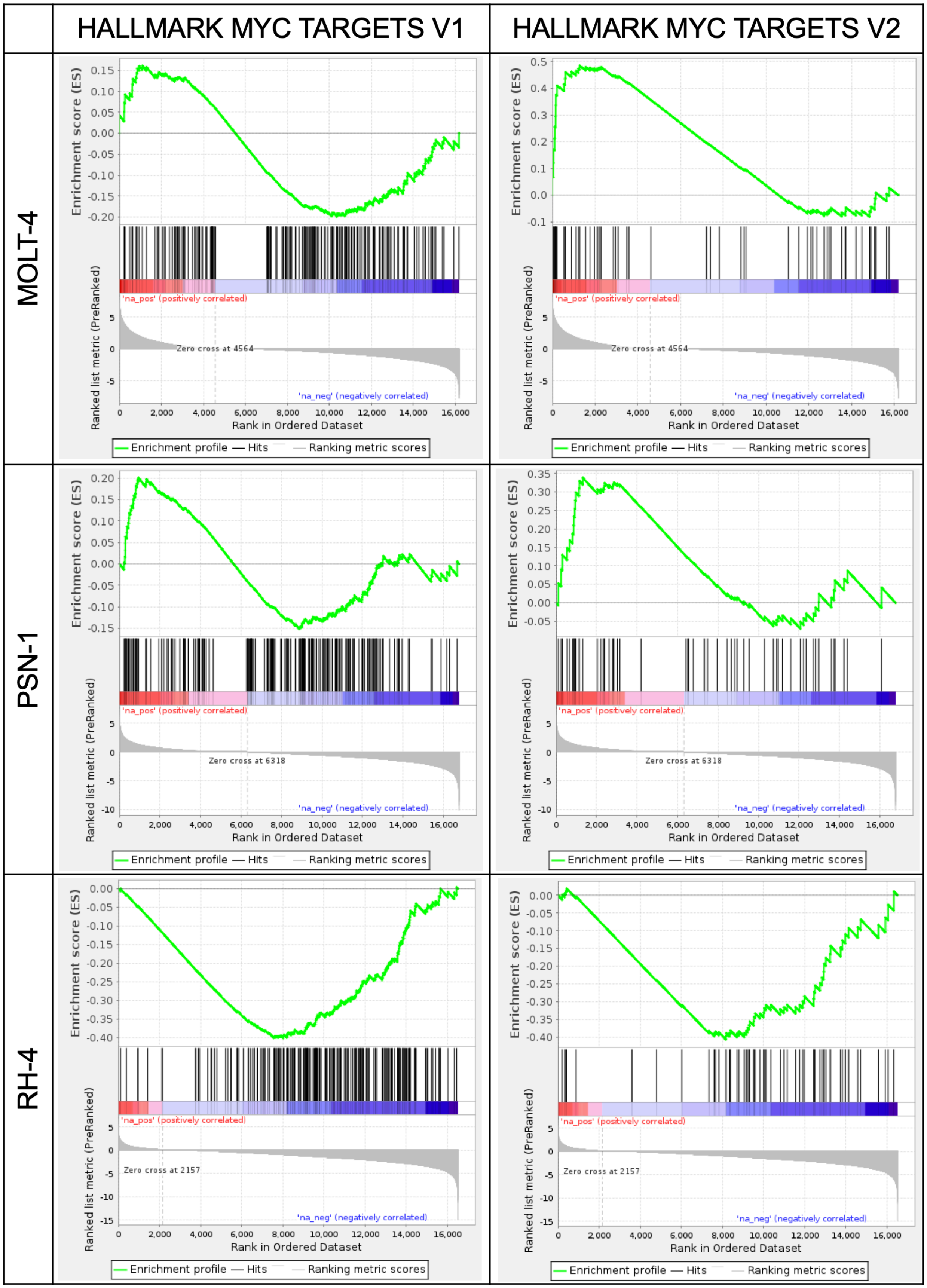


**Figure S3.2: Effects of CDK9 degradation vs inhibition on MYC targets across three cell lines**
Rank ordered gene set enrichment analysis resulting from RNA sequencing evaluation of transcript levels following 4 hours of treatment with 1.2 μM of inhibitor KB-0742 and 15nM of degrader KI-CDK9d-32 in MOLT-4, PSN-1, and RH-4. Differential expression was based on a direct comparison of inhibitor / degrader, thus positive correlates are indicative of degrader dominance in downregulating those transcripts.

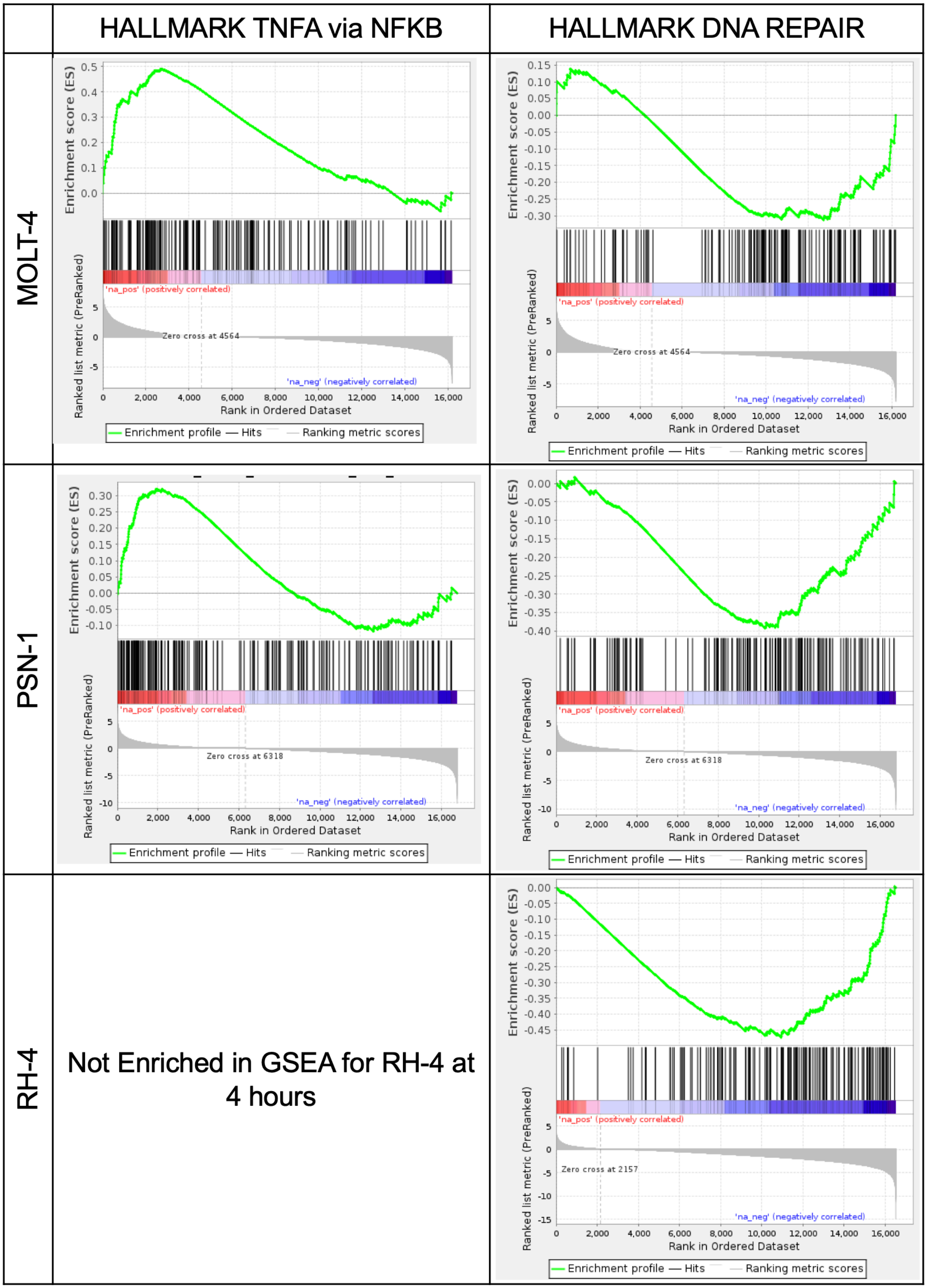


**Figure S3.3: Effects of CDK9 degradation vs inhibition on two key divergent gene sets**
Rank ordered gene set enrichment analysis resulting from RNA sequencing evaluation of transcript levels following 4 hours of treatment with 1.2 μM of inhibitor KB-0742 and 15nM of degrader KI-CDK9d-32 in MOLT-4, PSN-1, and RH-4. Differential expression was based on a direct comparison of inhibitor / degrader, thus positive correlates are indicative of degrader dominance in downregulating those transcripts.

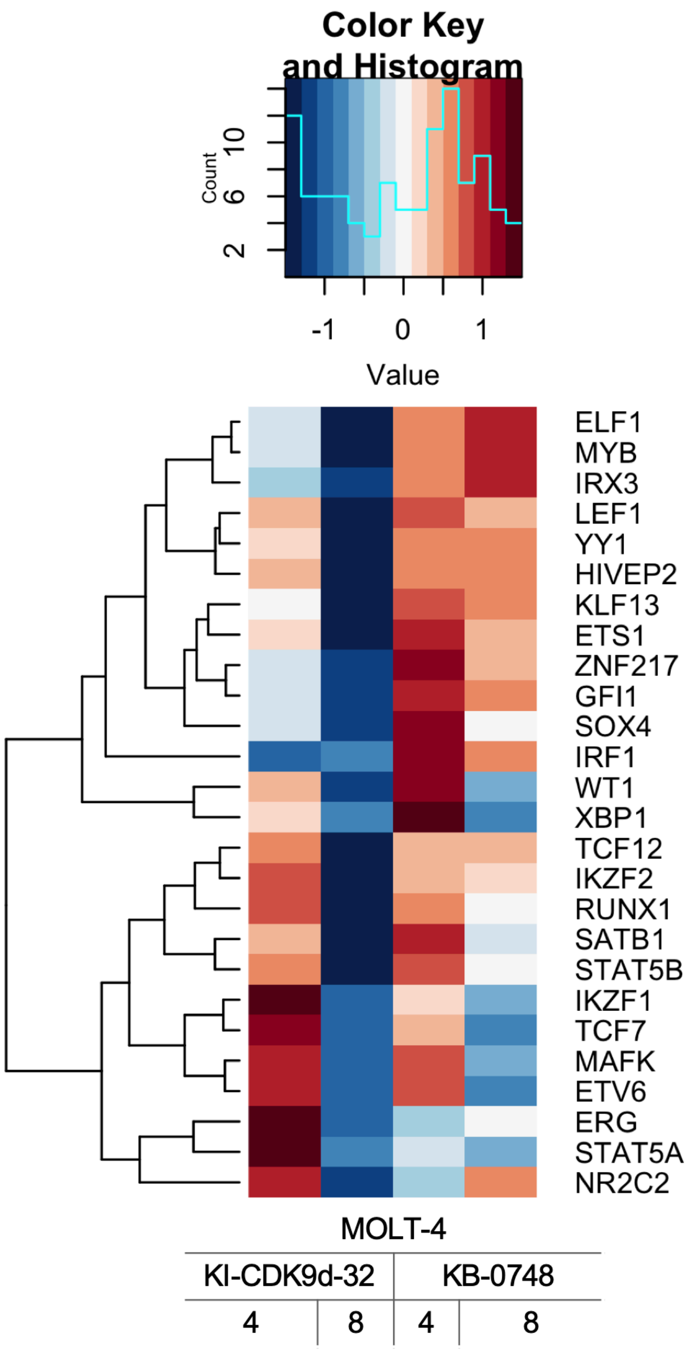


**Figure S3.4A: Effects of CDK9 degradation vs inhibition on core regulatory circuitry genes of MOLT-4**
RNA sequencing evaluation of transcript levels following 4, and 8 hours of treatment with 1.2 μM of inhibitor KB-0742 and 15nM of degrader KI-CDK9d-32 in MOLT-4. Differential expression was based on a direct comparison of inhibitor and degrader to DMSO treatment. Heatmap encodes the z-score log2 fold changes of significant genes ($p < 0.05$).

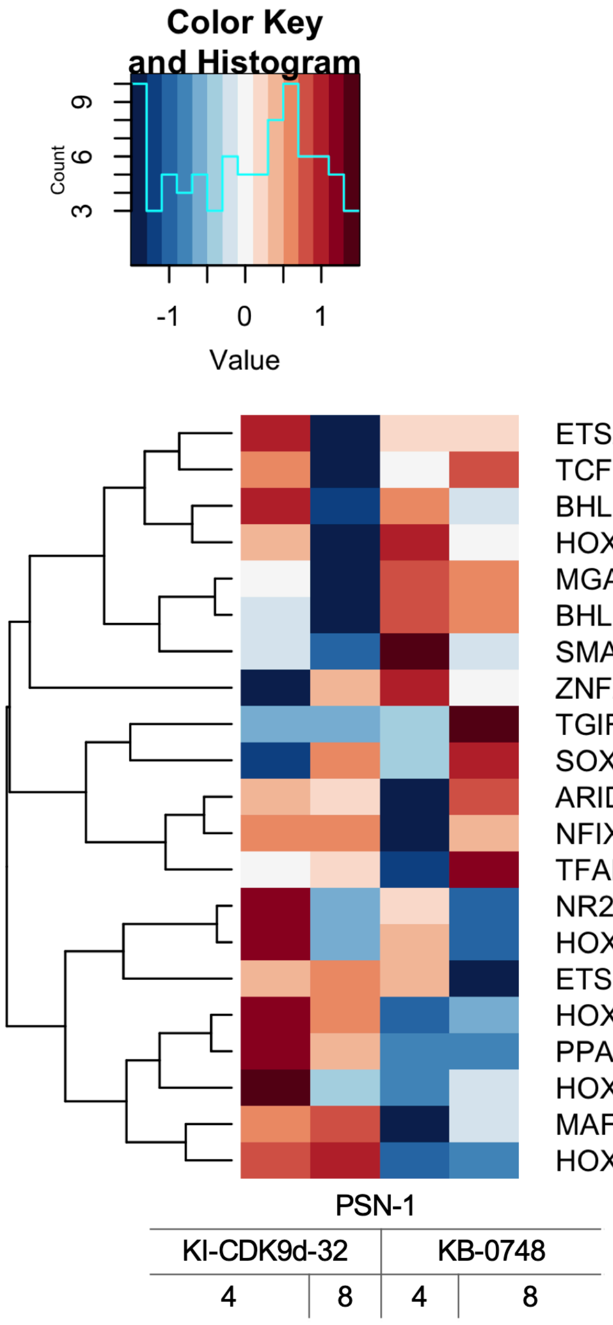


**Figure S3.4B: Effects of CDK9 degradation vs inhibition on core regulatory circuitry genes of PSN-1**

RNA sequencing evaluation of transcript levels following 4, and 8 hours of treatment with 1.2 μM of inhibitor KB-0742 and 15nM of degrader KI-CDK9d-32 in PSN-1. Differential expression was based on a direct comparison of inhibitor and degrader to DMSO treatment. Heatmap encodes the z-score log2 fold changes of significant genes ($p < 0.05$).

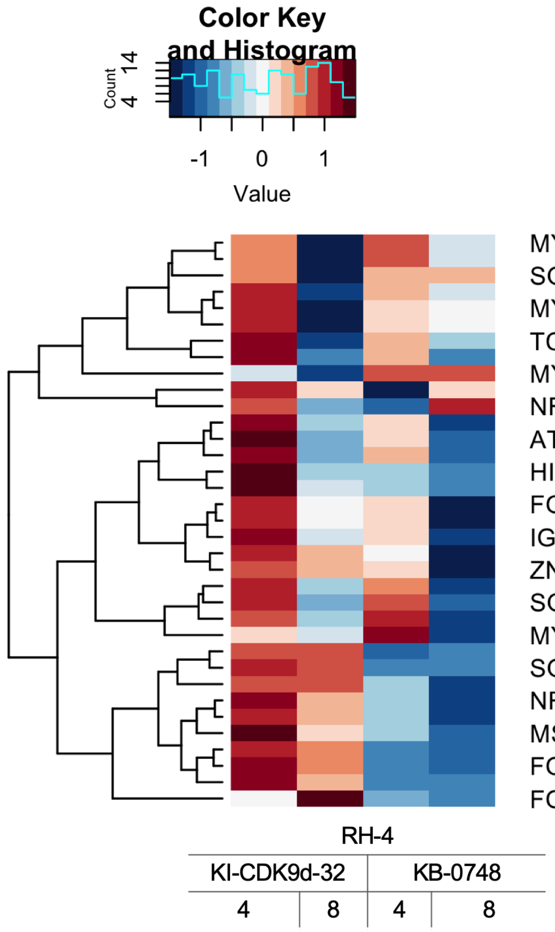


**Figure S3.4C: Effects of CDK9 degradation vs inhibition on core regulatory circuitry genes of RH-4**
RNA sequencing evaluation of transcript levels following 4, and 8 hours of treatment with 1.2 $\mu$M of inhibitor KB-0742 and 15nM of degrader KI-CDK9d-32 in RH-4. Differential expression was based on a direct comparison of inhibitor and degrader to DMSO treatment. Heatmap encodes the z-score log2 fold changes of significant genes ($p < 0.05$).

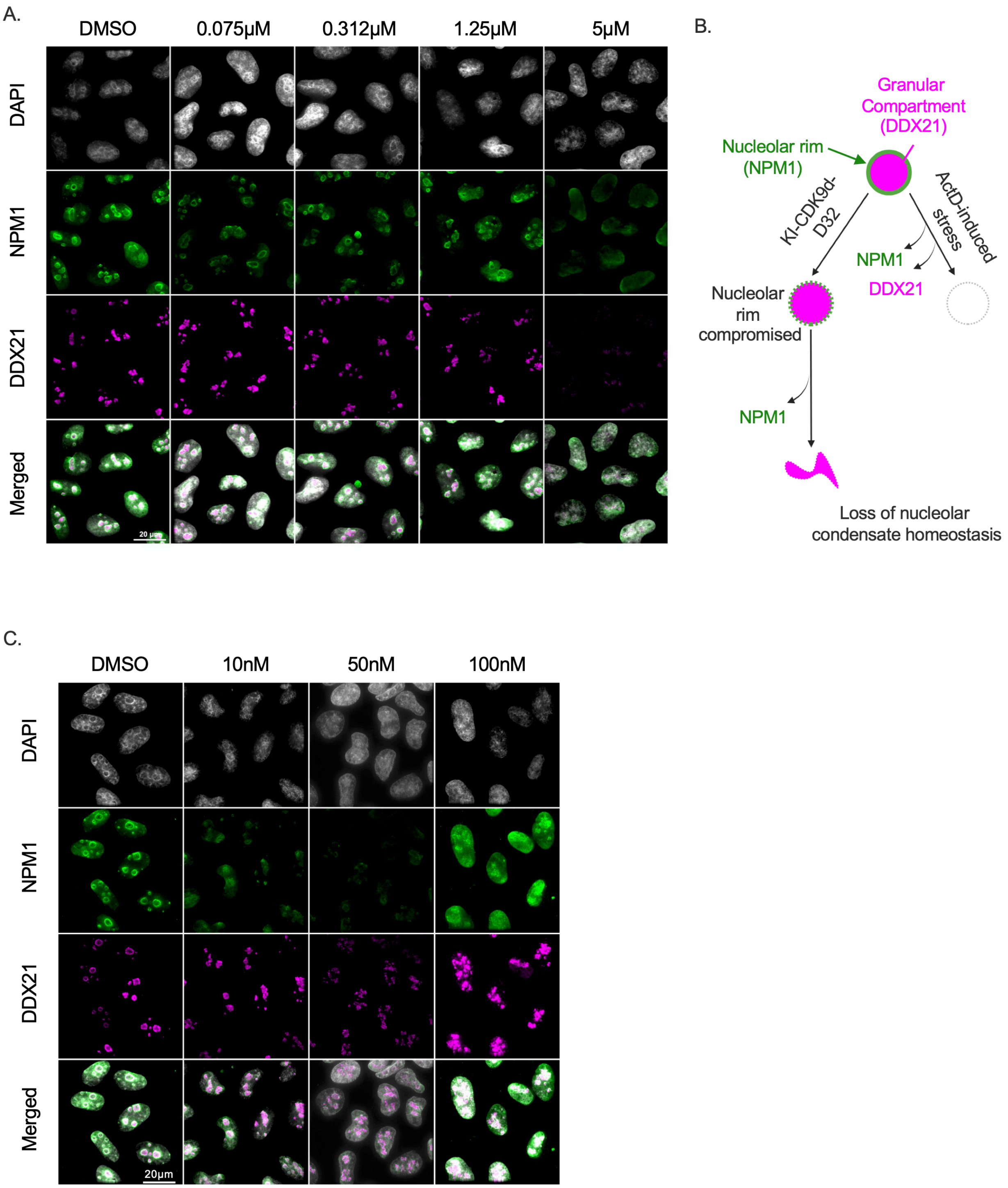


**Figure S4: KI-CDK9d-32 potently destabilizes nucleolar homeostasis**

A. Dose-response fluorescence imaging of HEK-293 cells following 4 hours of treatment with the inhibitor KB-0742, and DMSO control. The nucleolar rim is only compromised at the highest concentration of 5µM. B. Illustration of the nucleolar compartments. C. Dose-response fluorescence imaging of the effects of degrader KI-CDK9d-32 following 4 hours of treatment. Effects are visible at the lowest concentration tested. Images were processed using ImageJ.

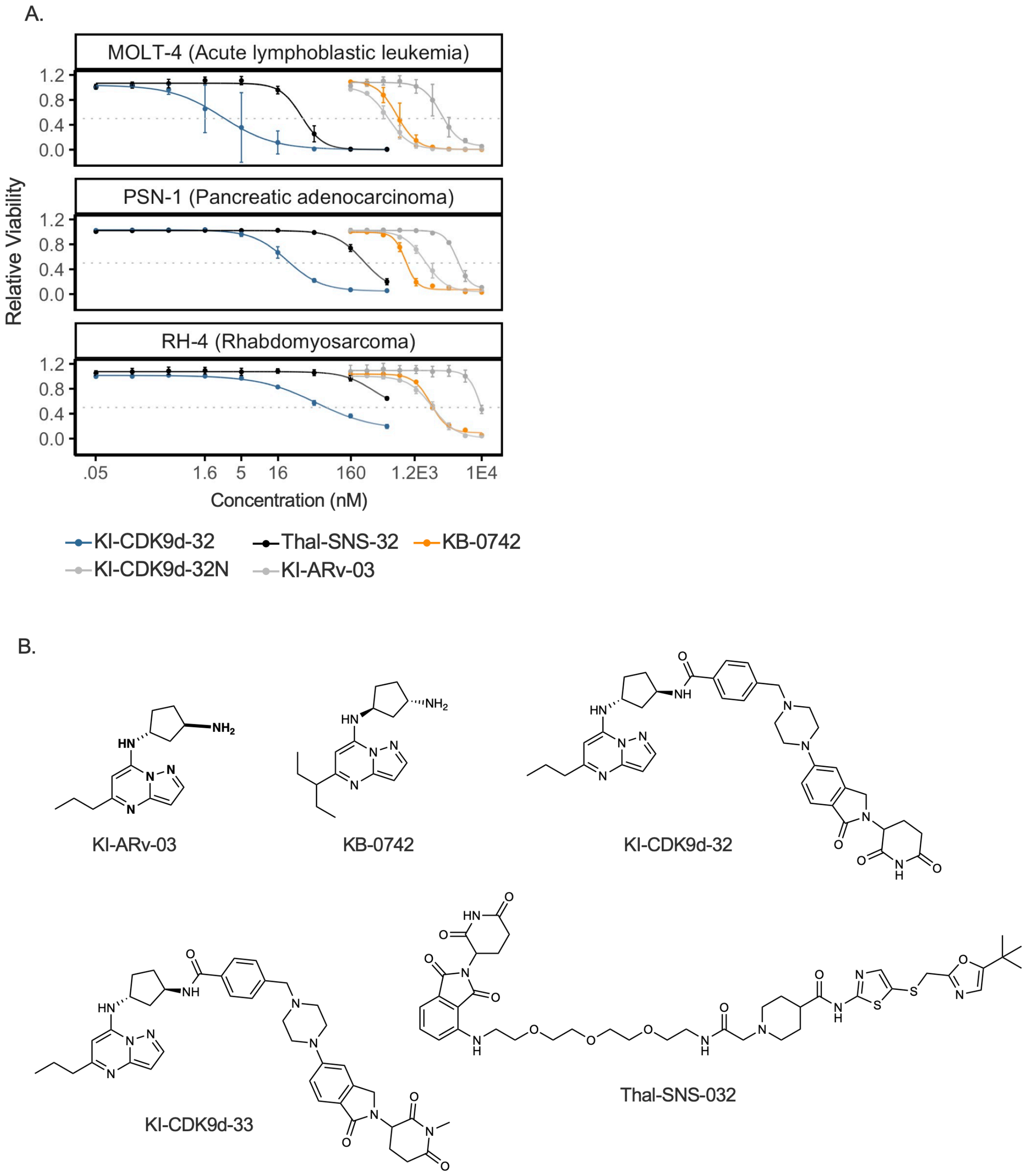


**Figure S5 : KI-CDK9d-32 demonstrates strong sensitivity in MOLT-4, PSN-1, and RH-4**

(A) Dose-response curves representing cell viability of three cell lines (MOLT-4, PSN-1, and RH-4) measured 72 hours post-treatment with a panel of compounds: KI-CDK9d-32, KI-CDK9d-32N, KI-ARv-03, KB-0742, and Thal-SNS-32. X-axis is log transformed with concentration shown in anti-log for readability. Error bars indicate the mean ± standard deviation from n=3 technical replicates. Concentration ranges for degraders were 0–500nM, while inhibitors and KI-CDK9d-32N were tested from 0–10µM. (B) Structures of compounds used in this experiment.

A.

| Compound ID | Assay Conc. (µM) | Mean Papp A-B (10-6 cm/s) | Mean Papp B-A (10-6 cm/s) | Mean (B-A/A-B) Efflux Ratio | Mean A-B % Recovery | Mean B-A % Recovery | A-B Permeability Ranking |
|---|---|---|---|---|---|---|---|
| KI-ARv-03-D08_NI_10uM | 10 | 0.00136 | 0.00397 | 2.92 | 0.772 | 0.799 | Lower |
| KI-CDK9d-32_NI_10uM | 10 | 0.0957 | 8.84 | 92.3 | 0.771 | 0.488 | Lower |
| Controls: | | | | | | | |
| Talinolol_NI_10uM | 10 | 0.156 | 5.43 | 34.9 | 0.947 | 0.862 | Lower |
| Talinolol_VERA_10uM | 10 | 0.314 | 2.75 | 8.74 | 0.955 | 0.864 | Lower |
| Ranitidine_NI_10uM | 10 | 0.357 | 1.31 | 3.65 | 0.999 | 0.841 | Lower |
| Warfarin_NI_10uM | 10 | 30.7 | 18.2 | 0.594 | 0.941 | 0.854 | Higher |

B.

| Compound ID | Conc (µM) | Matrix | Species | Matrix Conc (mg/mL) | Replicate | CLint (mL/min/kg) | CLint (L/h/kg) | Half Life (min) |
|---|---|---|---|---|---|---|---|---|
| KI-CDK9d-32 | 2 | Microsomes, Liver | Mouse, CD-1 | 0.5 | Average of 3 | 281 | 16.9 | 19.4 |
| Verapamil | 2 | Microsomes, Liver | Mouse, CD-1 | 0.5 | Average of 3 | 1410 | 84.6 | 3.86 |

**Table S1 : Assessment of Permeability and metabolic stability for KI-CDK9d-32**

A. This table presents the results of a Madin-Darby Canine Kidney (MDCK) cell line assay used to evaluate the apparent permeability coefficients (Papp) of KI-CDK9d-32 and four control compounds at a concentration of 10 µM. The assay measures directional permeability (A-B and B-A), efflux ratio, and recovery percentages to determine the compound's ability to permeate cell membranes, with higher Papp values indicating greater permeability. Compounds with an efflux ratio > 2 indicate a potential to be substrates for P-glycoprotein or other efflux transporters. The 'A-B Permeability Ranking' is determined based on the Papp A-B values, with a threshold of 1x10^-6 cm/s differentiating lower from higher permeability.

B. Summary of metabolic stability assessment in mouse liver microsomes. Verapamil is included as a reference compound. Higher CLint values and shorter half-lives indicate faster metabolism and lower stability of the compound. Both permeability and metabolic stability assessments were conducted by Charles River Laboratories following standard internal protocols.

Table S2

| Cluster 1 | Cluster 2 | Cluster 3 | Cluster 4 |
|---|---|---|---|
| OSRC2 | PATU8988T | HCC827 | NIHOVCAR3 |
| PANC1 | SKNMC | MCF7 | HEL |
| SCC25 | A673 | OCIAML5 | HEL9217 |
| HCC1937 | KARPAS299 | SKES1 | LS513 |
| KMRC20 | U87MG | PC3 | T24 |
| NCIH747 | NCIH2052 | G401 | PC14 |
| SNU449 | MFE280 | ZR751 | NCIH1650 |
| CAKI1 | DETROIT562 | GAMG | U118MG |
| TE11 | NCIH2122 | OCIAML2 | OV56 |
| NCIH1299 | G402 | NOMO1 | HUPT3 |
| K562 | J82 | CAL62 | OCILY19 |
| UACC257 | U937 | L363 | CFPAC1 |
| TE1 | NCIH1944 | CAL120 | THP1 |
| KMS11 | NCIH1792 | HUPT4 | SW1990 |
| HSC3 | C32 | CALU6 | RD |
| KYSE510 | KYM1 | KP4 | RCC10RGB |
| HCC15 | A2780 | BT549 | COLO320 |
| NCIH23 | LN18 | NB4 | SNU1079 |
| IPC298 | WM793 | 42MGBA | DAOY |
| COLO792 | RH30 | JURLMK1 | JHH6 |
| | HGC27 | RPMI7951 | A375 |
| | KYSE150 | JEKO1 | DEL |
| | CAL51 | MOLM13 | PANC0403 |
| | MFE296 | SUDHL4 | DANG |
| | AN3CA | RL | AU565 |
| | RL952 | COLO800 | KELLY |
| | SNGM | SAOS2 | SKNAS |
| | MONOMAC1 | TYKNU | TE10 |
| | OCUG1 | HUH7 | MSTO211H |
| | | KU1919 | MKN1 |
| | | CALU1 | MKN45 |
| | | A172 | SKHEP1 |
| | | RVH421 | U2OS |
| | | SW620 | HCC1143 |
| | | CAOV3 | OE33 |
| | | NCIH446 | TF1 |
| | | NB1 | LUDLU1 |
| | | MDAMB468 | HLF |
| | | HT | 769P |
| | | PF382 | SKMEL3 |
| | | NALM6 | UACC62 |
| | | 22RV1 | NCIN87 |
| | | IGROV1 | A704 |
| | | LNCAPCLONEFGC | MELHO |
| | | MFE319 | TE8 |
| | | 8305C | RT112 |
| | | 8505C | HUH1 |
| | | DOTC24510 | JHH4 |
| | | SKN | VMRCRCW |
| | | PFSK1 | SNU423 |
| | | | CORL88 |
| | | | CAL78 |
| | | | CAL33 |
| | | | KURAMOCHI |
| | | | ABC1 |
| | | | HT1197 |
| | | | IGR39 |
| | | | HT29 |
| | | | A498 |
| | | | EBC1 |
| | | | KMS27 |
| | | | NCIH1975 |
| | | | LN229 |
| | | | MIAPACA2 |
| | | | JHH1 |
| | | | KNS42 |
| | | | HCC1806 |
| | | | HEP3B217 |
| | | | HS944T |
| | | | NCIH441 |
| | | | 786O |

| | | | |
|---|---|---|---|
| | | | IGR37 |
| | | | SUIT2 |
| | | | CORL23 |
| | | | SBC5 |
| | | | SW1573 |
| | | | A549 |
| | | | KMRC1 |
| | | | OVCAR8 |
| | | | A3KAW |
| | | | HCC1395 |
| | | | TCCSUP |
| | | | HT1376 |
| | | | HEPG2 |
| | | | NCIH1703 |
| | | | SJSA1 |
| | | | NCIH196 |
| | | | MDAMB231 |
| | | | ONS76 |
| | | | KYSE30 |
| | | | CAMA1 |
| | | | A2058 |
| | | | SHP77 |
| | | | KATOIII |
| | | | BICR22 |
| | | | COLO679 |
| | | | KYSE410 |
| | | | SKMEL30 |
| | | | CAL27 |
| | | | FADU |
| | | | JHH7 |
| | | | HCC1954 |
| | | | NCIH358 |
| | | | AGS |
| | | | MELJUSO |
| | | | IGR1 |
| | | | SW1783 |
| | | | NCIH1793 |
| | | | SW1271 |
| | | | FTC133 |
| | | | MDAMB453 |
| | | | NUGC3 |
| | | | TE4 |
| | | | IM95 |
| | | | NCIH2172 |
| | | | SNU1 |
| | | | MDST8 |
| | | | RKO |
| | | | LS180 |
| | | | ISHIKAWAHERAKLIO02ER |
| | | | CCK81 |
| | | | DU145 |
| | | | HT115 |
| | | | MEWO |
| | | | JURKAT |
| | | | HCT15 |
| | | | SKMEL2 |
| | | | WM2664 |
| | | | CASKI |
| | | | SW756 |
| | | | HEC1 |
| | | | HT3 |
| | | | MCC13 |
| | | | MCC26 |
| | | | SISO |
| | | | CAL72 |
| | | | KMCH1 |

**Table S2 : Cell clusters based on sensitivity to KB-0742 and KI-CDK9d-32**

This table presents clusters of cells obtained following UMAP dimensionality reduction followed by HDBSCAN clustering of cell lines in both the PRISM pooled screen and the secondary individual cell line screen. The Z-score of IC50 and AUC values, relative metrics of compound sensitivity in the respective platforms, were used as inputs into UMAP to obtain a reduced dimensional data that used in HDBSCAN to obtain unbiased clustering of cell lines from the clusters obtained in Figure 6D.
